# Maternal ranging strategies facilitate offspring social play at energetic cost in the most solitary ape

**DOI:** 10.64898/2026.06.20.733430

**Authors:** Odd T. Jacobson, Alison M. Ashbury, Brendan J. Barrett, Margaret C. Crofoot, Paulina Kukofka, Julia A. Kunz, Sri Suci Utami Atmoko, Caroline Schuppli, Erin R Vogel, Carel P. van Schaik, Maria A. van Noordwijk

## Abstract

In most vertebrates, social play among peers is considered essential for behavioral development. Yet in solitary species bearing single offspring, opportunities for social play are inherently scarce. Whether mothers of such species actively facilitate play opportunities for their offspring, and at what cost, remains unknown. We used 15 years of behavioral and movement data (∼30,000 observation hours) from 31 wild Bornean orangutan (*Pongo pygmaeus wurmbii*) mother-offspring pairs to test whether mothers adjust ranging behavior to increase their offspring’s access to play with neighboring peers. Neighboring mothers with similarly aged offspring showed disproportionately high annual overlap in space use, independent of their relatedness or fruit availability. They intensified use of shared areas within existing range boundaries rather than shifting or expanding their ranges, indicating a fine-scaled ranging strategy. Mothers also incurred energetic costs; on days their offspring played with peers, mothers traveled farther and spent less time feeding. Travel distances were also elevated on the days before and after play, with mothers orienting movement toward play partners’ core areas before play and back toward their own core areas after play. This suggests these encounters are planned and actively pursued over multiple days rather than arising by chance. These findings reveal that orangutan mothers incorporate their infants’ social needs into daily ranging decisions, at a cost to their own energy budgets. This points toward an underappreciated form of maternal investment and illustrates how the social requirements of development can be met even near the solitary extreme of animal social organization.

## Introduction

Animal social organization varies along a spectrum, from cohesive groups with stable membership, to fission–fusion societies with fluid associations, to largely solitary lifestyles in which individuals forage and travel autonomously [1, 2]. Where a species falls along this continuum reflects an evolutionary balance of costs and benefits. Group living can reduce predation risk and improve access to information, mates, and cooperative partners, but it also intensifies resource competition and disease transmission [3, 4]. Species at the solitary end of the spectrum, for whom costs outweigh benefits, forgo many advantages of stable sociality. Yet solitary species are rarely truly asocial [5, 6]. They form selective social relationships, and their offspring share many of the same developmental needs as those of group-living species. Yet how these needs are met in solitary species, particularly those with single young and prolonged maternal dependence, remains poorly understood. Among these developmental demands, access to play partners may prove especially challenging. Play behavior is widespread across the animal kingdom, documented in both vertebrates (mammals, birds, reptiles, fish), and invertebrates (octopuses and insects) [7, 8]. Social play in particular, involving reciprocal play between peers, is thought to be especially important during development. It strengthens social skills and emotional regulation [9, 10], hones motor and competitive abilities [11, 12], and prepares juveniles to flexibly respond to unpredictable social and environmental challenges [13]. Early-life social play also predicts later survival and reproductive success in several wild mammal populations [14–17].

The developmental importance of social play poses a quandary for more solitary species. In group-living systems, access to play partners is often a routine by-product of social life. Solitary species that bear litters, including most felids and many other carnivorans, partially overcome this constraint through sibling play within the family unit [18]. But in many slow-reproducing solitary mammals (e.g., giant pandas [19], rhinoceroses [20], and orangutans [21]), offspring are typically reared as singletons, leaving them with limited access to siblings and other familiar peers. In such systems, opportunities for social play may depend on active maternal facilitation, potentially requiring mothers to alter their ranging, association patterns, and energy budgets to create social opportunities for their young. Whether mothers actually do so, and at what cost, remains unknown. Evidence for such behavior would constitute a substantial, previously overlooked form of maternal investment, revealing how the social requirements of development are met even near the solitary extreme of animal social organization.

Bornean orangutans offer a unique system to test whether mothers invest in social play opportunities for their offspring. They are the most solitary of the great apes, with adult females spending the majority of their time alone in the company of their dependent offspring [22]. Their reproductive pattern, characterized by single births and prolonged interbirth intervals averaging 7.6 years, typically results in mothers caring for only one immature offspring at any given time [21]. They are also among the largest-brained mammals [23]; given immature play scales with relative brain size [8], their need for play partners may be particularly pressing. Adult females occupy distinct but overlapping home ranges, such that every resident female has a unique set of neighbors [24, 25]. Despite large overlap [26], associations between neighboring mothers are rare, usually lasting only a few hours and seldom extending across consecutive days [27]. Mothers appear to gain little from these encounters. Unlike many group-living primates, female orangutans do not groom one another or form alliances with associates [26, 28]. Associations may even be costly, reducing feeding efficiency and increasing physiological stress [27, 29]. For immature offspring, however, these encounters provide their *only opportunities* to interact and play with peers— a potentially important context for cognitive development [30]. This raises the possibility that mother-mother associations are shaped by offspring rather than maternal needs.

Here, we leverage 15 years of behavioral and movement data (∼30,000 observation hours) from 31 wild Bornean orangutan (*Pongo pygmaeus wurmbii*) mother-infant pairs (each comprising a mother and her youngest, pre-weaned offspring) at Tuanan Research Station in Central Kalimantan, Indonesia (2*^◦^*09’S, 114*^◦^*26’E). This non-masting peat-swamp forest has few large, predictable fruit trees, making shared attraction to a food source an unlikely driver of social associations [31, 32]. We examine whether mothers adjust ranging behavior to increase their offspring’s access to play with neighboring peers, and whether they incur energetic costs while doing so. In this population, 11.9% of full follow days included some mother-mother association; and 6.31% included play between infants. Because orangutan births are not seasonally synchronized, neighboring infants naturally differ in age [33], providing the required variation to ask whether age similarity structures their play opportunities, as expected if play partners are generally matched in size and developmental stage. We apply Bayesian hierarchical Social Relations Models (SRMs), which account for the dyadic nature of neighboring mothers’ interactions, to establish whether infants play more with peers that are closer in age and assess whether mothers share space disproportionately with neighboring mothers whose infants are more similar in age to their own. We then use individual-level hierarchical models to examine whether play entails energetic costs to mothers by asking whether they travel farther and feed less on play days. Finally, to assess whether mothers actively pursue encounters rather than meet opportunistically, we examine whether increased travel extends to the days preceding and following play events, and whether mothers direct their movements toward the core areas of play partners before subsequently returning to their own.

## Results

### Similarly-aged infant dyads play more

We identified 266 unique infant dyads from 31 infants born to 14 mothers, observed across 79 mother dyads. Following our hypothesized causal structure (Figure S4), we fit a hurdle Poisson symmetrical SRM to annual play-scan counts per dyad, with joint observation effort (total behavioral scans during which both infants were alive and dependent) as an offset and fixed effects for offspring age difference, summed offspring age, maternal relatedness (*r*, ranging from 0 to 0.5), fruit availability (FAI), and vegetation greenness (EVI). The model included varying intercepts for mother dyad and for individual mothers via a multi-membership term (see Materials and Methods).

Offspring age difference had a credibly negative effect on play occurrence and frequency (*β* = –0.32 [-0.57,-0.09]; *PP >* 0 = 0.011; Figure 1a, b; model predictions over raw data in Figure S7)— infant pairs closer in age played together more often, adjusting for summed offspring age, maternal relatedness, and environmental conditions. The age-difference effect persisted (*β* = –0.32 [-0.57,-0.09]; *PP >* 0 = 0.011) when the model was offset by mother-mother association scans (i.e., when mothers were within ∼50 m) rather than total joint observation effort. This indicates that offspring themselves contribute to this pattern by playing more with similarly-aged peers once their mothers have brought them into proximity (Figure S9a; SI Results).

**Figure 1:**
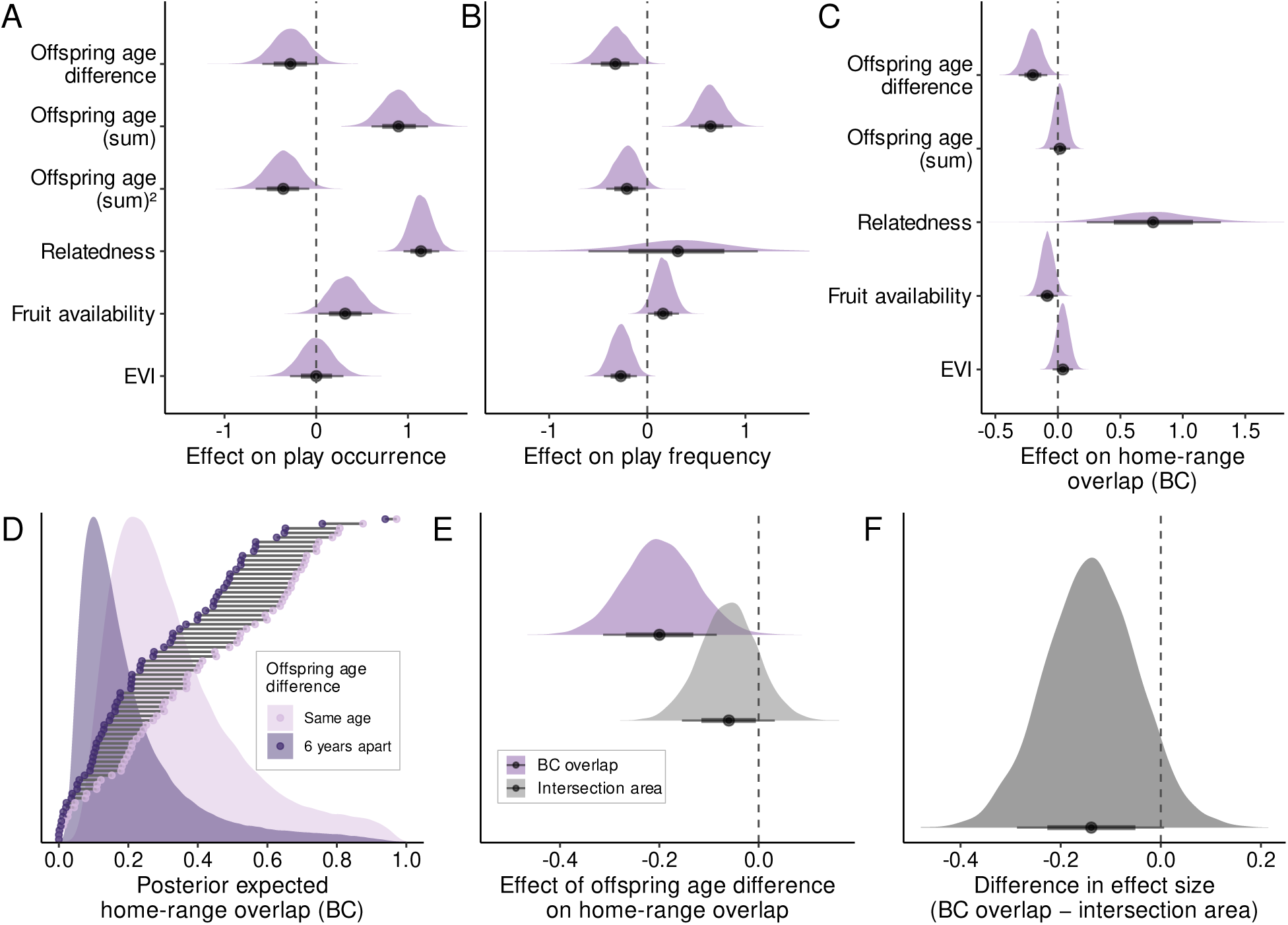
Mothers with similarly aged offspring share more space, and their offspring play more. Posterior effect sizes from social relations models (SRMs) of (a, b) infant-infant play scans (*n* = 996 infant-dyad-years across 79 mother dyads) and (c) mother-mother home-range overlap (Bhattacharyya coefficient; *n* = 282 mother-dyad years across 71 mother dyads). Panel (a) shows the hurdle component (probability of play occurrence), sign-flipped so positive values indicate higher probability of play occurrence; panel (b) shows the count component (play frequency). Points and thick and thin lines show posterior medians with 66% and 89% credible intervals, respectively. (d) Counterfactual home-range overlap for each dyad when offspring are the same age versus 6 years apart. Points show posterior medians across all mother-mother dyads (one horizontal bar per dyad); background densities show population-level predictions. (e) Offspring age-difference effect on home-range overlap versus intersection area. (f) Posterior difference between the two effects (BC overlap minus intersection area).

Summed offspring age showed a credibly positive linear effect with a negative quadratic component (linear *β* = 0.64 [0.43,0.85]; *PP >* 0 = 1; quadratic *β* = –0.21 [-0.42,-0.02]; *PP >* 0 = 0.04), indicating a developmental peak of play previously documented in this population [28]. Maternal relatedness strongly affected whether play occurred at all (*β_hurdle_* = 1.14 [0.96,1.34]; *PP >* 0 = 1; Figure 1a) but had no clear effect on how often it occurred once it did (*β* = 0.31 [-0.56,1.16]; *PP >* 0 = 0.728; Figure 1b). Both the occurrence and frequency of play increased with fruit availability (*β* = 0.16 [0,0.32]; *PP >* 0 = 0.952; Figure 1a,b).

### Mothers with similarly aged offspring use shared space more intensively

We estimated annual home ranges as autocorrelated kernel density estimates (AKDEs) from continuous-time movement models [34], which account for autocorrelation in GPS relocations and irregular sampling schedules. AKDEs produce utilization distributions (UDs), continuous probability surfaces describing the intensity of space use, from which we quantified pairwise overlap using the Bhattacharyya coefficient (BC; *BC* = 0.45 across 282 mother-dyad-years from 71 unique mother-mother dyads). Unlike geometric measures based on a single home-range contour, BC captures the shared intensity of space use across two individuals’

UDs on a scale from 0 (none) to 1 (identical). Following the same causal structure (Figure S4), we fit a Gaussian symmetric SRM on logit-transformed BC, incorporating measurement uncertainty from the AKDE confidence intervals, with fixed effects for offspring age difference, summed offspring age, maternal relatedness, FAI, EVI, and mean home-range area, and the same varying-effects structure as the play SRM.

Offspring age difference had a credibly negative effect on BC overlap (*β* = –0.2 [-0.32,-0.09]; *PP >* 0 = 0.003; Figure 1c): mothers whose offspring were closer in age used shared space more intensively (Figure 1d; Figure S6 for mapped examples), adjusting for summed offspring age, maternal relatedness, and environmental conditions. Maternal relatedness had a positive effect on overlap (*β* = 0.76 [0.24,1.31]; *PP >* 0 = 0.988), indicating that more closely related mothers shared space more.

To distinguish whether the age-difference pattern reflected more intensive use of shared space within existing range boundaries or shifts in range boundaries themselves, we refit the same model on intersection area (the geometric overlap of 95% UD contours). Offspring age difference had a more ambiguous effect on intersection area (*β* = –0.06 [-0.15,0.03]; *PP >* 0 = 0.144; Figure 1e), and the posterior difference between the two effect sizes indicated that the offspring age-difference effect operated primarily through fine-scaled adjustments in space use within overlapping ranges rather than through shifts in range boundaries (Figure 1f). A complementary analysis of behavioral association data showed a similar tendency: mothers with more similarly-aged offspring tended to associated more in space and time (Figure S9b).

### Mothers travel farther and feed less on days their offspring play

We estimated daily path length (DPL) for 14 mothers across 2211 days using the continuous-time speed and distance (CTSD) method [35], which accounts for irregular GPS sampling and location error. We quantified daily feeding effort from 2-minute instantaneous scan sampling, calculating the proportion of scans spent feeding across 2347 days for 14 mothers. We fit separate Bayesian Generalized Linear Multilevel Models (GLMMs) — a Gamma model for DPL incorporating measurement uncertainty derived from CTSD confidence intervals, and a beta-binomial model for feeding scans out of total daily scans — each conditioning on offspring age, fruit availability, EVI, and including varying intercepts per mother.

Because social association can elevate maternal energetic costs through resource competition [29], we used a three-level categorical predictor distinguishing days on which the focal mother was alone with her offspring (no association, no play; reference), days on which she associated with another neighboring conspecific without infant play (association, no play), and days on which infant-infant play occurred (association, play). This allowed us to separate the cost of play itself from the cost of association. Mothers travelled farther and fed less on days they associated with neighboring conspecifics, even when no infant play occurred (*β_DP L_* = 0.18 [0.14,0.22]; *PP >* 0 = 1; *β_feed_*= –0.15 [-0.19,-0.11]; *PP >* 0 = 0; Figure 2). On play days, mothers traveled even farther and fed less compared to association-only days (*β_DP L_* = 0.34 [0.28,0.41]; *PP >* 0 = 1; *β_feed_* = –0.25 [-0.32,-0.18]; *PP >* 0 = 0). The posterior difference between play days and association-only days was positive for daily path length (Δ = 0.16 [0.08,0.23]; *PP >* 0 = 1) and negative for feeding (Δ = –0.1 [-0.18,-0.03]; *PP >* 0 = 0.015), suggesting an energetic cost on play days beyond what association alone explains (SI Results).

**Figure 2:**
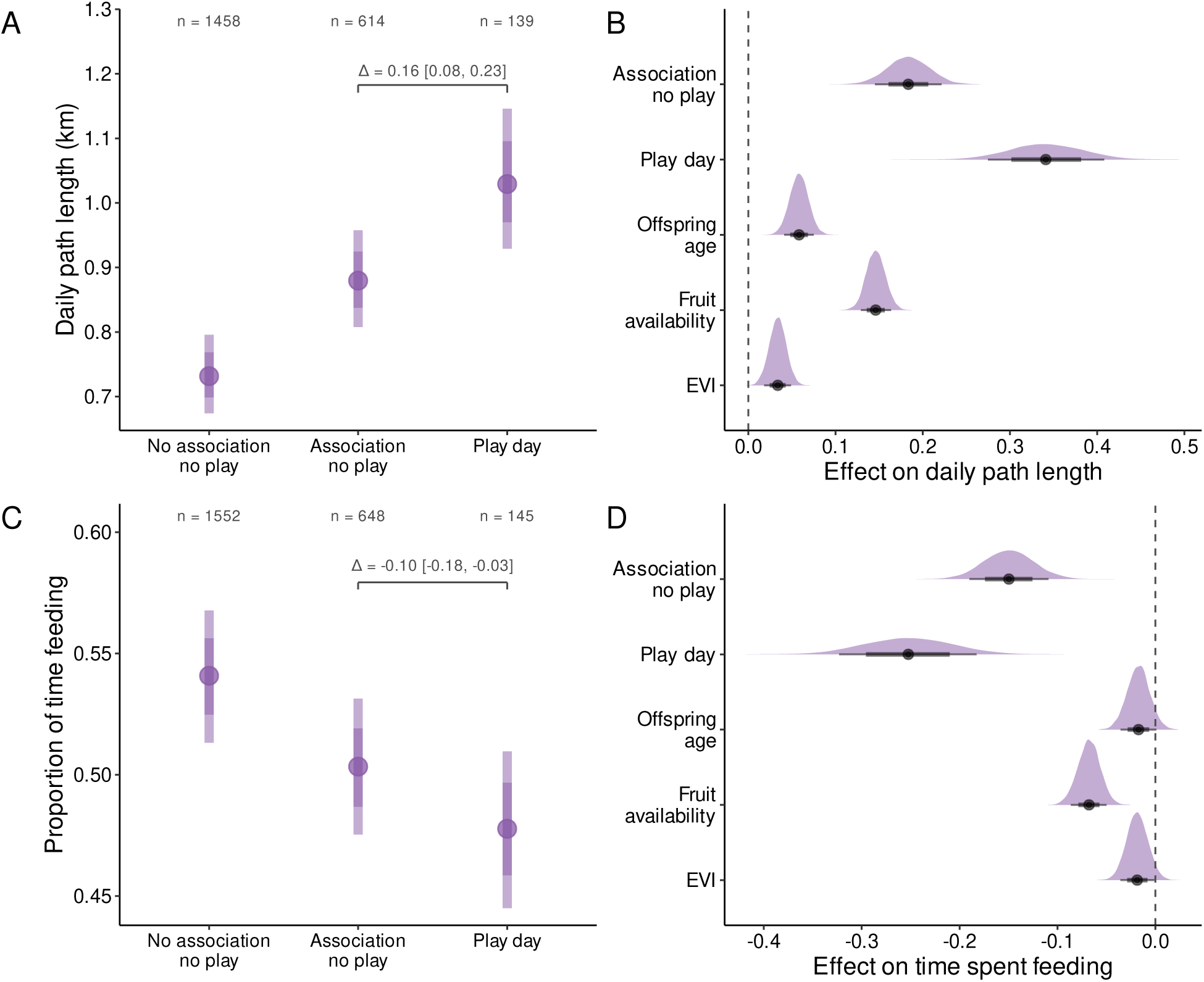
Mothers travel farther and feed less on days their offspring play. Posterior predictions and effect sizes from Bayesian GLMMs of daily path length (*a, b*) and proportion of time feeding (*c, d*), comparing days when the focal mother was alone with her offspring (no association, no play; reference), days when she associated with another neighboring conspecific without infant play (association, no play), and days when infant-infant play occurred (play day; which by definition involves association). (*a, c*) Posterior predicted values across the three categories, with covariates held at their mean values; sample sizes (*n*) are shown for each category. Sample sizes differ between panels because path length estimation requires sufficient GPS fixes while feeding estimation requires behavioral observation scans; not all days met both criteria. Points and dark and light lines show posterior medians with 66% and 89% credible intervals, respectively. The bracket and labelled value within each panel indicate the posterior median and 89% credible interval of the difference between play days and association-only days, isolating the additional cost of play beyond what association alone produces. (*b, d*) Posterior effect sizes for all covariates; densities show the full posterior distributions, and points with thick and thin lines show posterior medians with 66% and 89% credible intervals.

### Mothers travel toward and then away from play partners across multiple days

To assess whether mothers’ behavioral adjustments were confined to play days or extended over multiple days, we classified each day as either a non-play day (reference category), the day before, the day of, or the day after an offspring play event, restricting the analysis to days for which adjacent-day classification could be determined reliably (SI Materials and Methods). We fit the same Bayesian GLMM structure as above, replacing the three-level categorical play predictor with this four-level categorical variable. Mothers traveled farthest on play days themselves (*β* = 0.34 [0.27,0.41]; *PP >* 0 = 1; Figure 3a), but also tended to travel farther on the day before (*β* = 0.09 [-0.02,0.21]; *PP >* 0 = 0.904) and the day after (*β* = 0.14 [0.05,0.23]; *PP >* 0 = 0.992) play events compared to non-play days. In contrast, the proportion of time spent feeding was lower only on play days (*β* = –0.23 [-0.31,-0.16]; *PP >* 0 = 0; Figure 3b), with no clear difference on days before (*β* = 0.03 [-0.09,0.15]; *PP >* 0 = 0.675) or after (*β* = –0.04 [-0.14,0.06]; *PP >* 0 = 0.279).

**Figure 3:**
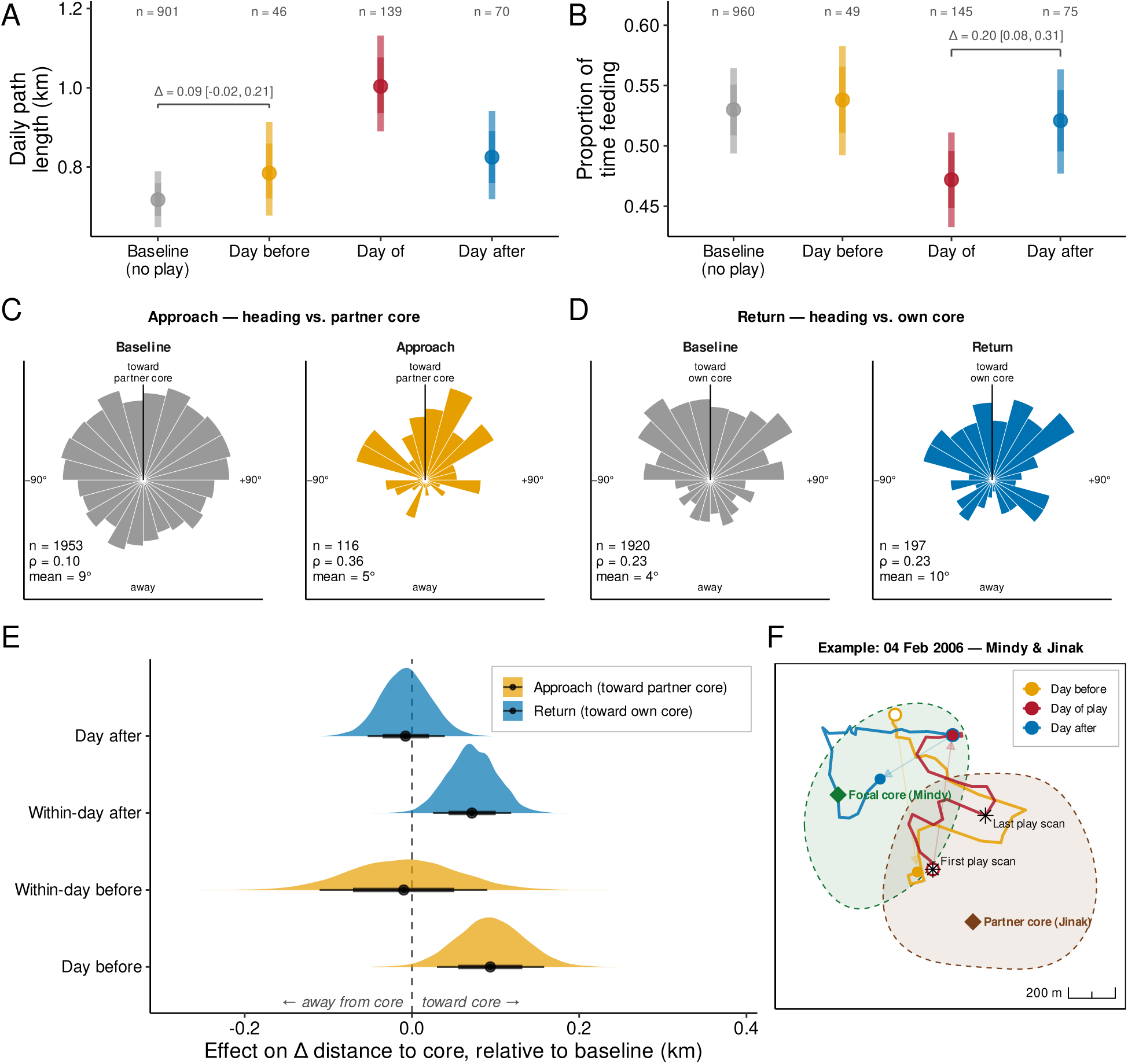
Mothers travel toward and then away from play partners across multiple days. (a, b) Posterior predicted values of daily path length and proportion of time feeding across play adjacency categories, with covariates held at their mean values. Points and dark and light lines show posterior medians with 66% and 89% credible intervals, respectively. Within each panel, the labelled bracket indicates the posterior median and 89% credible interval for a single representative contrast (a: day before vs. no play; b: day of vs. day after); contrasts between other adjacent categories can be inferred visually from their relative positions. (c, d) Distribution of focal-day net travel direction relative to the reference core (toward partner’s core in c, toward focal’s own core in d) for the Approach (c) and Return (d) phases. Bins are 15*^◦^* wide; per-phase mean direction, mean resultant length *ρ* (ranging from 0 = uniform to 1 = all directions identical), and sample size are shown. Sample sizes differ across panels because the four analyses (daily path length, feeding, Approach, Return) apply different retention criteria; see SI Materials and Methods for details. (e) Posterior effect sizes on Δ distance to goal across four temporal phases relative to baseline, from Bayesian regressions controlling for relatedness, core separation, offspring ages, fruit and EVI availability, daily path length, and observation window duration, with focal and dyad random intercepts. Thick and thin lines show 66% and 89% credible intervals, receptively. (f) Example dyad-day showing one mother’s daily paths the day before, day of, and day after her offspring plays with neighboring peer. Open circles mark each day’s first fix, filled circles the last, joined by arrows indicating overall daily displacement; dashed polygons show each mother’s current-year 50% utilisation distribution; diamonds mark the prior-365-day mean location of each mother (reference core); stars mark the first and last observed play scans.

To distinguish whether mothers’ elevated travel around play days reflected directed movement toward partners, we tested two complementary models: 1. an Approach model (do mothers move closer to the partner’s range core before play?) and 2. a Return model (do mothers move back toward their own range core after play?). Mothers’ travel was credibly oriented toward the partner’s core on the day before play (*β* = 0.09 [0.03,0.16]; *PP >* 0 = 0.991; Figure 3c,e); no clear directional bias was detected in the pre-play within-day window (*β* = –0.01 [-0.11,0.09]; *PP >* 0 = 0.439). After play, mothers redirected travel back toward their own core within the same day (*β* = 0.07 [0.03,0.12]; *PP >* 0 = 0.994; Figure 3d-f), with no remaining directional shift relative to baseline by the day after (*β* = –0.01 [-0.05,0.04]; *PP >* 0 = 0.392). Circular distributions supported this pattern: headings were strongly concentrated toward the partner’s core during Approach phases, but only weakly oriented toward the mother’s own core during Return phases, relative to an already homeward-oriented baseline (Figure 3c,d).

## Discussion

Our study provides strong evidence that wild Bornean orangutan mothers adjust their ranging behavior to increase their offspring’s access to social play. Mothers whose offspring were closer in age shared space more intensively, and their offspring played more, while mothers incurred measurable short-term energetic costs through increased travel and reduced feeding on play days. Greater travel distance and directional orientation around play events further suggest that these encounters arise from multi-day adjustments in maternal movement rather than chance encounters. Together, these findings illustrate how offspring social needs can shape maternal ranging decisions, even when those decisions impose immediate energetic costs.

How do mothers achieve these encounters? Our findings indicate that mothers do so through finer-scale adjustments in how they use space, by concentrating activity in areas already shared with particular neighbors rather than shifting or expanding overall range boundaries. This supports previous findings showing that related and unrelated females overlap similarly in range boundaries, whereas related females associate far more frequently within those shared areas [26]. Our results suggest that offspring age similarity drives the same pattern: mothers whose infants are closer in age concentrate their activity in shared areas, increasing the probability of encounter and thus of offspring play. Crucially, offspring age similarity predicted both maternal space sharing and offspring play even after accounting for maternal relatedness. Nonetheless, offspring of females with no known related neighbors were rarely seen in peer play, suggesting that some baseline of kin-based familiarity is a prerequisite for peer play in this population.

Greater travel distance around play events, together with directional movement toward play partners before play and back toward mothers’ own core areas afterward, further suggests that these encounters emerge from deliberate, multi-day adjustments rather than opportunistic co-occurrence. Such behavior is consistent with evidence from Tuanan that female associations are flexible and responsive to social and ecological context [24, 31]. Moreover, these patterns point toward the advanced spatial cognition and route planning documented in orangutans [36, 37], and with broader evidence for future-oriented planning in birds and mammals [38]. Orangutan mothers’ ranging thus reveals decisions that integrate offspring social needs with broader ecological priorities across both space and time.

This pattern contrasts with mechanisms facilitating offspring play in other species where social contact is infrequent or episodic. Brown bear cubs from different families play when mothers converge at concentrated food resources such as salmon streams, making play a by-product of maternal foraging rather than targeted social facilitation [39, 40]. Among many fission-fusion species such as spider monkeys, offspring play occurs during maternal associations maintained for direct affiliative benefits, including grooming and alliance support [41]. Meanwhile, giraffe calves are cared for in communal nursery groups (crèches), where peer social opportunities arise naturally while mothers forage elsewhere [42]. Across these systems, offspring gain social access in contexts where mothers also appear to receive independent benefits from association. In contrast, our results suggest that orangutan mothers gain no direct return: they travel farther, feed less, and adjust their space use in ways best explained by the developmental needs of their offspring. These costs also exceed those attributable to association alone [27]. We propose two non-exclusive mechanisms: maternal vigilance while infants are with playmates likely distracts mothers from feeding, whereas locating and remaining near parties containing suitable play partners imposes travel costs beyond the scramble competition inherent to association [29].

That orangutan mothers incur these costs despite little immediate return is particularly striking given their extreme energetic constraints [32, 43], presumably exacerbated by their arboreality. Orangutans have among the lowest metabolic rates of any mammal [44] and the longest interbirth intervals of any terrestrial animal [21]. So, even modest changes in daily expenditures may carry meaningful consequences for maternal energy balance and reproductive timing. Consistent with this, offspring social play increased with fruit availability, suggesting that play is more readily afforded when ecological constraints are relaxed (though peer-play did not cease entirely during periods of scarcity [31]). The costs we document here therefore represent an underappreciated form of maternal investment [45, 46] that extends beyond carrying, provisioning, and protection. Orangutan mothers are known to support offspring development through prolonged parental care and opportunities for social learning [47–50]. Our results reveal an additional aspect of this investment—the active shaping of an offspring’s social environment.

Comparable patterns are observed in some large-scale human societies, where parents invest considerable time and effort managing children’s peer access by arranging social contacts and restructuring daily routines (e.g., [51–53]; cf. [54] for small-scale societies). Similarly, in fission-fusion chimpanzees, mothers may alter infants’ early social exposure by adjusting their own gregariousness, despite the costs of association (e.g., competition, infanticide risk) [55]. Such investment may yield delayed fitness returns. For example, in feral horses, greater maternal investment predicts more juvenile play, better body condition, and higher survival [14]. In brown bears and bottlenose dolphins, juveniles that play more survive longer or reproduce more successfully [15, 17]. If similar benefits accrue in orangutans, the short-term energetic costs borne by mothers may be offset by long-term gains through improved offspring competence, survival, or future reproduction.

Our findings extend the growing recognition that solitary mammals can be socially more complex than once assumed [5, 6]. Recent work has emphasized that species lacking stable groups can nonetheless maintain selective and enduring adult relationships [56]. Our results add a new dimension to this social complexity. Mothers appear to adjust movement and space use to locate suitable play partners for their infants, navigating a social landscape where opportunities are intermittent and depend on the developmental state of both their own offspring and their neighbors’. Such behavior requires combining knowledge of where neighboring females range with knowledge of their offspring’s ages, consistent with the idea that facultative sociality (e.g., semi-solitary and fission-fusion systems) places distinctive demands on social cognition by requiring individuals to track relationships and anticipate opportunities across extended spatial and temporal scales [57]. This holds especially where mothers cannot depend on reliable chance meetings at key resources. More broadly, the challenge of solitary living may lie not in reduced social developmental needs, but in the parental effort required to meet them. Such systems may thus preserve key social experiences for offspring while shifting the associated energetic and cognitive costs onto mothers.

Our results reveal a robust behavioral pattern consistent with active maternal facilitation of offspring play. Play events coincided with systematic shifts in maternal movement and activity budgets, including longer travel and movement oriented toward partner core areas before play. These findings support the interpretation that mothers plan and actively structure social opportunities for their offspring. Future work using route-choice or state-space approaches could test this more directly by distinguishing social-seeking from foraging-associated movement behaviors (e.g., area-restricted search) [58, 59]. The next challenge is to determine whether these short-term maternal costs are offset by longer-term gains, including improved post-weaning offspring survival or reproductive success. Only long-term field studies can determine whether the hidden maternal costs documented here are ultimately repaid through the success of the next generation.

Orangutans are often described as the most solitary of the great apes, yet our findings show that mothers in this species incorporate offspring social needs into daily ranging decisions at a cost to their own energy budgets. This represents an overlooked form of maternal investment, extending beyond offspring nutrition and protection to include their social development. Such investment in social play opportunities arises in a species where ecological pressures otherwise favor minimal sociality, suggesting that the need for social experience during development is a deep constraint on mammalian life histories. Across the spectrum of social organization, species may meet these shared developmental needs through fundamentally different social strategies. If similar patterns occur more broadly, offspring social development may represent both an underappreciated benefit of group-living and a largely hidden cost of solitary living.

## Methods

### Study site and data collection

We analyzed long-term data from wild Bornean orangutans at the Tuanan Research Station, Central Kalimantan, Indonesia (2003-2018). Trained observers conducted nest-to-nest focal follows of identified mothers and their pre-weaned offspring (the youngest immature still ranging with the mother, in all cases under nine years), logging GPS locations at ∼30-min intervals and recording 2-min instantaneous scans of activity (including feeding and social play) and association partners throughout the day [26]. The dataset comprised 32631.1 observation hours across 3448 focal follow days, including 2211 full-day nest-to-nest follows. Across full follow days, infant-infant play was typically absent, averaging only 2.37 scans per day (median = 0). We derived offspring ages from long-term birth and demographic records [21], and computed a continuous coefficient of maternal relatedness (*r*) for each dyad from long-term maternal pedigrees [60] (SI Materials and Methods).

### Ethics statement

This observational study of wild orangutans involved no experimental manipulation or interaction with focal individuals. Observers maintained distances of at least 10 m whenever possible, and data collection complied with Indonesian legal requirements under permits approved by RISTEK-BRIN, KSDAE-KLHK, and the Ministry of Internal Affairs.

### Environmental covariates

We characterized food availability using a monthly fruit availability index (FAI) derived from long-term phenological monitoring of marked trees [32]. As a landscape-scale index of canopy greenness and vegetation productivity variation (including the effects of drought and fire), we extracted monthly Enhanced Vegetation Index (EVI) values from MODIS MOD13Q1 Collection 6.1 within a 6 × 6 km window centered on Tuanan [61, 62]. Full details are provided in SI Materials and Methods.

### Movement and space-use metrics

We estimated annual home ranges, pairwise overlap, and daily path length using continuous-time movement models in ctmm, which account for location error, temporal autocorrelation, and irregular GPS sampling [34]. Annual home-range analyses used GPS locations from both full-day and partial-day follows, provided they met predefined criteria for temporal spread and number of independent GPS locations (see SI Materials and Methods) [63]. Daily path length analyses were restricted to full-day nest-to-nest follows. For each mother-year (a single mother’s movement data over one calendar year), we estimated autocorrelated kernel density home ranges and quantified pairwise overlap as the Bhattacharyya coefficient (BC) between utilization distributions [64, 65]. We estimated daily path length with the continuous-time speed and distance method [35]. Uncertainty in BC and daily path length estimates was propagated into downstream models as measurement error.

### Statistical analyses

We used directed acyclic graphs (DAGs) to define target effects and identify adjustment sets for all statistical models; full DAGs, statistical notation, and reasoning behind covariate choices are provided in SI Materials and Methods.

### Dyad-level statistical analyses

We analyzed three dyadic outcomes — pairwise home-range overlap (BC), geometric home-range overlap (intersection area), and infant-infant play frequency — using hierarchical symmetric (or undirected) social relations models (SRMs) [66–69]. SRMs partition dyadic variation into individual-level contributions (e.g., each mother’s general tendency to overlap with) and a dyad-specific component (the unique pairwise deviation from those individual tendencies), enabling efficient partial pooling across repeated dyads [70]. Every model included varying intercepts for mother-dyad identity and for individual mothers via a multi-membership term [71].

We modeled annual BC overlap on the logit scale as a Gaussian outcome, incorporating the 95% confidence interval around each estimate as measurement error so that CTMM-derived uncertainty propagated into the posterior. We modeled annual intersection area (km^2^) with a hurdle-Gamma likelihood and log link. Both home-range models included standardized fixed effects for the absolute offspring age difference within the dyad, the summed offspring ages, maternal relatedness, monthly fruit availability index (FAI), mean EVI over the sampling window, and the mean home-range area of the two mothers (to partial out baseline differences in space-use extent). We included FAI and EVI as covariates throughout to account for environmental variation in food availability and habitat productivity over time, improving the precision of main effects (e.g., offspring age difference).

We modeled annual infant-infant play scans with a hurdle-Poisson SRM, with the log of observation effort across the two infants during their joint dependency window (total scans when both were alive and immature) as an offset. The play model included the same fixed effects as the overlap models (offspring age difference, summed offspring age, maternal relatedness, FAI, and EVI), with the exception of mean home-range area, and additionally a quadratic term in summed offspring age to capture the non-monotonic developmental trajectory of play previously documented in this population [28]. The home-range models were fit at the mother-dyad-year level, whereas the play model was fit at the infant-dyad-year level but retained the same mother-dyad SRM structure to account for shared mothers and repeated dyads.

### Individual-level statistical analyses

#### Daily travel distance and feeding time

We tested whether maternal movement and feeding effort differed on days when the focal mother’s infant was observed engaging in social play, using four Bayesian generalized linear multilevel models (GLMMs). For each outcome (daily path length and feeding proportion) we fit two complementary specifications. The first used a three-level predictor contrasting days on which the focal mother was alone with her offspring (no play, no association; reference), days on which she associated with another neighboring conspecific without infant play (no play, association), and days on which infant-infant play occurred (which by definition involve association). We classified each focal day according to whether play or association occurred: a day was scored as containing play or association when at least one 2-min instantaneous scan recorded infant-infant play or association with a neighboring conspecific, respectively. We used binary daily occurrence rather than duration because our hypothesis concerned whether play or association occurred, not whether longer bouts imposed proportionally greater costs. Of 139 play days, 134 (96%) involved a matriline relative as one of the focal mother’s associates, consistent with kin-biased female association in this population [26]. Of 614 association-only days, 270 (44%) involved a matriline relative; the remainder involved adult males, unknown individuals, or unrelated females.

We modeled daily path length (km) with a Gamma likelihood and log link, incorporating the 95% confidence interval around each estimate as measurement error so that CTMM-derived uncertainty propagated into the posterior. We modeled daily feeding proportion with a beta-binomial likelihood (logit link) on counts of feeding scans out of total scans per follow day. All models included focal infant age, FAI, and mean EVI as covariates (all standardized), and a varying intercept for mother identity. DPL and feeding proportion are coupled both behaviorally (mothers allocate time within a 24-hour day) and energetically (faster patch depletion increases travel; more travel raises both resource encounter rate and caloric demand); we modeled them separately because they capture partly independent aspects of mothers’ daily strategies (Figure S14c). After conditioning on covariates, the residual correlation between the two outcomes was small (median *r* = −0.06, 89% HDI [−0.09, −0.04]; Figure S14b).

#### Directional travel relative to play partners

To test whether elevated travel around play days reflected directed movement, we measured whether mothers moved closer to or farther from a relevant range core over each travel window. For the *Approach* model, the relevant core was the play partner’s range core; for the *Return* model, it was the focal mother’s own range core. Range cores were defined as the mean of each mother’s GPS fixes over the 365 days preceding, and excluding, the focal play date (see SI Material and Methods for further detail). We calculated change in distance to the relevant core as the starting distance minus the ending distance, so that positive values indicate movement toward the core. Within play days, we also calculated pre-play and post-play windows, defined as the periods before the first observed play scan and after the last observed play scan, respectively. The *Approach* model used change in distance to the partner’s core as the response and contrasted non-play days (baseline), the day before play, and the within-day pre-play window. The *Return* model used change in distance to the mother’s own core as the response and contrasted baseline, the within-day post-play window, and the day after play. Both models included relatedness, core separation, offspring age, fruit availability, EVI, daily path length, and observation-window duration as covariates, with random intercepts for focal mother and mother-mother dyad. For descriptive directional analyses, we also calculated each mother’s net movement bearing and the bearing from her starting location to the relevant core. We summarized the angular difference between these bearings to assess whether movements were oriented toward the relevant core, aggregating the day-before and within-day pre-play phases for Approach and the within-day post-play and day-after phases for Return (Figure 3c,d; SI Materials and Methods).

### Model fitting and inference

We fit all Bayesian models in brms with cmdstanr [71]. We used weakly informative priors and assessed convergence with *R*^^^, effective sample sizes, posterior predictive checks (Figure S15), and trace plots. We summarized posteriors with medians, 89% highest-density intervals, and the posterior probability that each effect was greater than zero (*PP >* 0); see SI for summaries and diagnostics.

## Data Availability Statement

All analysis code and the data needed to reproduce the statistical models, figures, and in-text results are openly available on Edmond [72]. Reproducing the preprocessing steps additionally requires the raw activity and GPS location data, which are restricted under existing data-sharing agreements and to protect the whereabouts of critically endangered orangutans. Location data are archived under limited access on Movebank, and are available on reasonable request.

## Supporting information

Full Supplementary Information

## Acknowledgements

We thank the Indonesian State Ministry for Research and Technology and the National Research and Innovation Agency (RISTEK-BRIN), the Indonesian Institute of Sciences (LIPI-BRIN), the Directorate General of Natural Resources and Ecosystem Conservation-Ministry of Environment and Forestry of Indonesia (KSDAE-KLHK), the Ministry of Internal Affairs, the local government of Central Kalimantan, the Nature Conservation Agency of Central Kalimantan (BKSDA), the Kapuas-Kahayan Forest Protection Management Unit (KPHL Kapuas-Kahayan), the Bornean Orangutan Survival Foundation (BOSF), and BOS-MAWAS in Palangkaraya for permission to conduct research at Tuanan and for long-term support. We thank Universitas Nasional (UNAS), especially Dr. Tatang Mitra Setia, for long-standing collaboration with the Tuanan Orangutan Research Project since 2003. We are grateful to all who contributed to behavioral data collection and data management, especially Abuk, Idun, Isman, Pak Rahmat, Tono, Suwi, and Ibu Fitriah. This work was supported by the University of Zurich, the A. H. Schultz Foundation, the Swiss National Science Foundation (grant nos. 310030B_160363/1 and 3100A-116848), and the Max Planck Institute of Animal Behavior. This work was also supported by the Alexander von Humboldt-Stiftung through funds awarded to M.C.C. Open access funding was provided by the Max Planck Institute of Animal Behavior.

## Notes

https://doi.org/10.17617/3.XDGZMM

### Competing Interest Statement

The authors have declared no competing interest.

