## Supplementary material for "Maternal ranging strategies facilitate offspring social play at energetic cost in the most solitary ape": Full Supplementary Information

June 19, 2026

#### Materials and Methods

##### Estimating pairwise relatedness from matriline pedigrees

We assigned a coefficient of maternal relatedness ($r$) to each female-female dyad in the analysis ($n = 71$ dyads, 282 dyad-years, among 14 mothers; Figure 1) using maternal pedigrees compiled from long-term demographic records of the Tuanan Orangutan Research Project. Following standard pedigree-based coefficients [1], mother-daughter dyads were assigned $r = 0.5$ ($n = 6$); maternal half-sisters ($n = 3$) and grandmother-granddaughter dyads ($n = 2$) were assigned $r = 0.25$; aunt-niece dyads $r = 0.125$ ($n = 4$); and maternal cousins $r = 0.0625$ ($n = 1$). Females with no documented common maternal ancestor in the pedigree were assigned $r = 0$ ($n = 55$).

We restricted our relatedness measure to the maternal line. Although paternities are available for a subset of Tuanan individuals [2], what matters for the behavioral questions examined here is *knowable maternal relatedness*: the kin a female has had life-long social exposure to through her mother. Female orangutans are strongly philopatric and remain near their natal range, whereas males disperse widely and as adults range and reproduce far beyond their birthplace [3–5]. A female's maternal kin therefore form the stable social environment in which she develops, while her paternal half-sibs are typically scattered across the landscape and cannot be recognized through maternal social bonds. All values correspond to demographically observed mother-offspring links in the long-term records, with no values inferred or estimated.

##### Environmental covariate processing

We characterized local food availability using the fruit availability index (FAI), derived from long-term phenological monitoring at Tuanan, in which trained researchers recorded the percent of marked trees bearing fruit along permanent transects [6, 7]. FAI provides a direct monthly estimate of fruit abundance within the study area and was matched to each observation window by month (Figure 11).

We complemented FAI with the Enhanced Vegetation Index (EVI), a satellite-derived proxy for canopy greenness designed to remain sensitive in dense tropical canopies where NDVI often saturates [8]. We extracted EVI from the MODIS MOD13Q1 Collection 6.1 product, which provides 250 m, 16-day composite imagery. Using the `MODISTools` R package [9], we retrieved all pixels within a $6 \times 6$ km window centered on Tuanan Research Station. We averaged EVI values across pixels within each 16-day composite and then across composites within each month. Because satellite vegetation indices cannot resolve specific phenophases such as fruiting [10], we interpret EVI as an index of broad-scale leaf cover and drought-driven productivity variation rather than food abundance per se (Figure 12).

#### Movement data processing and space-use estimation

We estimated annual home ranges, pairwise home-range overlap, and daily path length (DPL) using the `ctmm` R package [11]. Continuous-time movement models account for temporal autocorrelation, irregular GPS sampling, and location error, all of which are common in nest-to-nest focal follows with handheld GPS devices. We specified a user-equivalent range error (UERE) corresponding to location errors of approximately 2-29 m, consistent with handheld GPS performance under tropical forest canopy. Location data used for annual home-range estimation included both full-day and partial-day follows, whereas DPL analyses were restricted to full-day follows, defined as follows beginning at morning nest departure and ending at evening nest arrival.

Before estimating annual home ranges, we excluded mother-years with clearly inadequate sampling coverage by requiring at least 8 observation days, 3 unique weeks, and 50 GPS locations. For each retained mother-year, we fitted candidate CTMMs and selected the best-supported model using $AIC_c$. We then estimated annual home ranges as autocorrelated kernel density estimates (AKDEs) [12] with optimal weights to reduce bias from uneven sampling [13]. We retained only home ranges with acceptable diagnostics, based on visual inspection of empirical and model-predicted variograms and an effective sample size for area $\geq 3$. This yielded 99 annual home ranges from 14 mothers with sufficient annual spatial coverage.

We quantified pairwise home-range similarity as the Bhattacharyya coefficient (BC) between utilization distributions using `ctmm::overlap()`. This procedure propagates uncertainty from the underlying CTMM fits into 95% confidence intervals on BC, which we carried into downstream Bayesian models as measurement error on the response. We estimated DPL using the continuous-time speed and distance (CTSD) method [14]. For each full-day follow, we fitted a per-day CTMM and multiplied the estimated mean speed by the total daily sampling duration. This approach is robust to variation in sampling rate, path tortuosity, and location error. For each day, we extracted the CTSD point estimate and its 95% confidence interval, which we propagated into downstream Bayesian models as measurement error on the response. This yielded 2211 daily path lengths from 14 mothers with eligible full-day follows. The sample sizes differed between annual home-range and daily path-length analyses because the two metrics required different forms of sampling coverage.

#### Play adjacency classifications and directed-travel methods

##### Play adjacency classifications

Each follow day was classified relative to play events as a play day (day-of), the day immediately before or after a play day (day-before, day-after), or a non-adjacent day (baseline). Play days took precedence over adjacent-day classifications, so the day-before and day-after categories capture only non-play days flanking play events. Days that fell within one day of play events on both sides (e.g., a non-play day with play occurring both yesterday and tomorrow) were excluded from the day-before and day-after categories to avoid ambiguity in their classification. These classifications are used in the daily path length, feeding, and directed-travel adjacency analyses.

##### Range core estimation

For each mother on each focal date, we defined her range core as the mean location of all GPS fixes collected during the preceding 365 days, excluding the focal date itself. We used this GPS-fix mean, rather than an AKDE-based core, because fitting a separate rolling AKDE for every mother-date combination was computationally impractical. To ensure stable estimates, we retained only (mother, date) pairs with $\geq$100 fixes spanning $\geq$3 unique weeks across $\geq$2 unique months within the 365-day window.

##### Travel vectors and within-day windows

For each candidate day, we computed the focal mother's net travel vector from her first GPS fix to her last fix of the day. For within-day windows on play days, we measured the focal mother's net travel vector from the first fix of the day to the first observed play scan (pre-play), and from the last play scan to the last fix (post-play). Within-day windows were retained only if they contained ≥3 fixes spanning ≥60 minutes with ≥30 m of net displacement.

##### Directional and distance measures

We then computed two quantities per record: (i) the angular difference between the net travel bearing and the bearing from the starting location to the reference core (partner's core for the Approach analysis; the focal mother's own core for the Return analysis), used as input to descriptive circular statistics; and (ii) the change in net distance to the reference core (distance from start to core minus distance from end to core, with positive values indicating net travel toward the core), used as the response variable in the Bayesian regressions described in Other dyadic and individual-level models.

##### Descriptive circular statistics

For each record we computed the angular difference between the focal mother's net travel bearing and the bearing from her starting location to the reference core. Day-before and within-day pre-play observations were pooled for the Approach, and within-day post-play and day-after observations were pooled for the Return. Per-phase summaries are the mean direction and mean resultant length $\rho$, ranging from 0 (uniform) to 1 (all observed directions identical), computed using the `circular` R package [15].

##### Sample size differences across analyses

The daily path length, feeding time, Approach, and Return analyses all draw from the same pool of nest-to-nest follow days but apply different retention criteria. Daily path length requires successful CTSD path-length estimation; feeding requires available behavioural scan data; missingness differs slightly between the two. The Approach and Return analyses additionally require a qualifying 365-day prior centroid for both mothers (see above), ≥50 m net daily displacement, ≥100 m starting distance to the reference core, and ≥200 m mode-separation between cores. These thresholds are required so that net travel direction can be meaningfully resolved. The Approach and Return baselines exceed those of the daily path length and feeding analyses because the dyad-indexed structure assigns one baseline row per focal-date per potential play partner (defined as any other mother whose infant was observed playing with the focal's infant at some point during the study; see Other dyadic and individual-level models). The rose-plot categories in Figure 3c-d also combine full-day adjacent observations with within-day pre- and post-play windows, exceeding the day-before/day-after counts in Figure 3a-b.

#### Causal inference framework and DAG specification

We framed each analysis as a question about causal effects rather than associations, and used directed acyclic graphs (DAGs) to make the assumed causal structure of the system explicit. A DAG encodes which variables we assume cause which others and lets us identify the minimal set of covariates needed to estimate a target causal effect without bias [16–18]. We constructed our DAGs using the `dagitty` package [19] in `R`. We treat each DAG as an explicit, falsifiable set of assumptions about the system, drawn from biological reasoning rather than learned from the data.

Throughout this section we use a set of terms from the causal inference literature. The *exposure* is the variable whose effect we want to estimate (e.g., offspring age difference). The *outcome* is the variable that effect acts on (e.g., home-range overlap). An *estimand* is the specific causal quantity we aim to estimate, defined by an exposure, an outcome, and the variables held fixed. A *confounder* is a variable that causes both the exposure and the outcome and would bias the estimate if not adjusted for. A *mediator* is a variable through which part of the exposure's effect on the outcome is transmitted. A *backdoor path* is any non-causal path between the exposure and outcome that would carry spurious association if left open.

Our statistical models fall into two causal families. The dyadic models (home-range overlap and infant–infant play) share an exposure (offspring age difference within a mother–mother dyad) and a confounding structure (Figure 4). The individual-level models (daily path length and feeding time) share an exposure and a distinct confounding structure (Figure 5). We present one DAG per family.

##### Estimands

For the overlap models (BC overlap vs. intersection area), the estimand is the direct causal effect of offspring age difference on dyadic home-range overlap within a dyad-year. The two models share an estimand and a DAG and differ only in the outcome scale. For the infant–infant play model, the estimand is the *total* causal effect of offspring age difference on play. We estimate the total rather than direct effect because our key question is whether dyads with similar-aged offspring play more overall, integrating both the direct mechanism (infants of similar age preferring to play once encountered) and the indirect mechanism (mothers of similar-aged infants overlapping spatially and creating encounter opportunities). For the day-level models, the estimand is the direct causal effect of focal-day offspring play (i.e., "play day") on the focal's daily energetic outcome (daily path length and feeding time).

##### DAG for the dyadic models

The exposure is offspring age difference (OFF_AGE_DIFF); the outcomes are home-range overlap (OVER) and infant–infant play (PLAY). Observed covariates are mother–mother relatedness (REL), mean dyadic home-range area (HRA), combined offspring age (OFF_AGE_SUM), combined mother age (MAGE), fruit availability (FAI), and vegetation greenness (EVI). Latent nodes capture a landscape-level environmental state (ENV), an unobserved spatial food-distribution variable (DISTRIB), and residual unmeasured drivers of past reproductive timing ($U_{\text{OFF\_AGE\_DIFF}}$, e.g., maternal body condition and prior infant survival). The full set of arrows is shown in Figure 4.

We highlight five assumptions. First, REL → OFF_AGE_DIFF: kin dyads have non-random offspring-age structure as a demographic consequence of relatedness, since mother–daughter dyads are necessarily separated by at least one interbirth interval and sister dyads by their mother's interbirth interval, while unrelated dyads face no such constraint. This makes REL a confounder of the OFF_AGE_DIFF → OVER relationship and required in the adjustment set. Second, REL → OVER, PLAY: kin share space and tolerate proximity at higher rates [3]. Third, OVER → PLAY: greater spatial overlap creates more opportunity for infant–infant encounter, making OVER a mediator on one of the paths from OFF_AGE_DIFF to PLAY. Fourth, MAGE causes OFF_AGE_DIFF, OFF_AGE_SUM, and HRA: a young mother cannot have an offspring age gap larger than her own reproductive history allows, mothers with longer reproductive histories tend to have older offspring, and older mothers occupy more established home ranges. This makes MAGE a confounder of the OFF_AGE_DIFF → OVER, PLAY relationships. Fifth, OFF_AGE_SUM and OFF_AGE_DIFF share unmeasured causes ($U_{\text{OFF\_AGE\_DIFF}}$, MAGE) but neither causes the other; both are functions of offspring birth dates rather than of each other. The remaining assumptions are as follows: ENV drives FAI, EVI, and DISTRIB; productivity and food distribution shape HRA; HRA, FAI, and EVI contribute to OVER; OFF_AGE_SUM contributes to PLAY directly and to HRA; and OFF_AGE_DIFF acts on OVER and PLAY.

##### Adjustment

For the direct effect of OFF_AGE_DIFF on OVER, there are two minimal sufficient adjustment sets: {EVI, FAI, HRA, REL} or {MAGE, OFF_AGE_SUM, REL}. These are equivalent for identification: backdoor paths can be closed either by intercepting environmental and ranging effects at HRA or by intercepting demographic effects upstream at MAGE and OFF_AGE_SUM. We adopt the first set for the overlap model and additionally condition on OFF_AGE_SUM for precision; none of these are descendants of OFF_AGE_DIFF, so this does not introduce post-treatment bias. For the total effect of OFF_AGE_DIFF on PLAY, the only minimal sufficient adjustment set is {MAGE, OFF_AGE_SUM, REL}. Unlike the overlap model, no alternative set based on HRA exists, because HRA does not lie on any backdoor path to PLAY in our DAG. The play model conditions on OFF_AGE_SUM (entered with a quadratic term to capture nonlinearity in developmental stage) and REL, and additionally on FAI and EVI for precision. We do not include MAGE in the primary models for two reasons (see *Maternal age* below).

##### Maternal age

The DAG identifies MAGE as a confounder of OFF_AGE_DIFF → (OVER), (PLAY). We do not include it as a fixed effect in the primary models for two reasons. First, stable between-mother variation in mother age is fully absorbed by the multi-membership random intercept on mother identity. The remaining unhandled component is within-mother aging across the study period, which is uniform across mothers (all aged ∼15 years together over 2003–2018) and is largely captured by OFF_AGE_SUM, which tracks the same temporal axis. Second, mothers' birth years are uncertain: eleven of fourteen mothers were already adults when observation began in 2003. Their birth years were reconstructed by working backward from the youngest known offspring using estimated interbirth intervals and age at first reproduction, yielding minimum-age estimates that underestimate true age by an unknown individual-specific amount (older sons that dispersed before observation began, infants that did not survive, daughters that emigrated to areas outside the study). To verify robustness, we report sensitivity analyses (Figure 8) in which mother age is included via a measurement-error specification [20] that propagates encoding uncertainty into the posterior. The focal offspring age difference effect remained directionally consistent across all specifications.

##### DAG for the day-level models

The exposure is offspring social play on the focal day (PLAY). The outcomes are daily path length (DPL) and the proportion of scans spent feeding (FEED). Observed covariates are the focal mother's offspring age (OFF_AGE), mother age (MAGE), FAI, and EVI; ENV is again the latent parent of the observed environmental proxies (Figure 5).

The arrows we assume are: OFF_AGE → PLAY (offspring engage in social play differently across developmental stages); OFF_AGE → DPL, FEED (maternal travel and feeding patterns vary with offspring developmental stage, as mothers adjust the pace of travel to their offspring's locomotor capacity and adjust feeding effort to changing energetic demands of dependence); MAGE → OFF_AGE (younger mothers have younger offspring), and MAGE → PLAY, DPL, FEED (mother age may independently affect activity budgets, ranging, and social behavior); FAI, EVI → PLAY, DPL, FEED (productivity and phenology shape both the opportunity for play and the travel and feeding required to meet daily energetic needs); FAI, EVI, DISTRIB → DPL (food distribution shapes how far mothers must travel to access resources); and PLAY → DPL, FEED (mothers monitoring offspring as they play takes time away from feeding which could have downstream effects on daily travel; mothers may also have to travel further to find suitable play partners).

##### Adjustment

The minimal sufficient adjustment set for the direct effect of PLAY on either outcome is {EVI, FAI, MAGE, OFF_AGE}. We condition on OFF_AGE, FAI, and EVI directly. We verify in sensitivity analyses (Figure 8) that explicitly including MAGE as a measurement-error covariate does not qualitatively alter the results.

##### Assumptions and limitations

For the dyadic models, our DAG cannot rule out the possibility of unmeasured time-varying confounders that affect both reproductive timing and ranging behavior. We also assume FAI and EVI together adequately proxy the latent environmental state; because they capture distinct aspects of productivity (ground-based fruit vs. remote-sensed canopy greenness) and are not collinear in our data; we consider this defensible but imperfect.

For the day-level models, plausible unmeasured day-level confounders include weather, chance encounters with neighbouring mothers, and within-day maternal state. We further assume the PLAY → DPL arrow runs in the direction we have drawn it, that mothers' elevated travel reflects pursuit of play opportunities rather than play being an incidental by-product of travel undertaken for other reasons.

##### DAG considerations for the directed-travel models

The Approach and Return models analyze a day-level exposure (play-adjacent phase) and an individual-level response (the focal mother's change in distance to the reference core — the partner's core for Approach, the focal's own core for Return). Because each row relates to a specific (focal, partner) dyad, both models use the same adjustment set as the dyadic models (relatedness, summed offspring age, offspring age difference, FAI, and EVI; Figure 4). Mode separation (distance between focal and partner cores), daily path length, and observation-window duration are included to capture geometric and sampling features that likely affect approach and return distances but are not part of the DAG.

#### Statistical formulation of the dyadic home-range overlap model

We modelled the annual pairwise home-range overlap between mothers as a symmetric Social Relations Model (SRM) [21–23], in which the dyadic outcome is decomposed into individual-level contributions and a dyad-specific component. Because our home-range overlap metric is symmetric by construction (the overlap between mothers $i$ and $j$ is identical to the overlap between $j$ and $i$), we used a symmetric SRM in which both members of a dyad contribute through a single shared random effect on individual identity, implemented as a multi-membership term [20]. This contrasts with asymmetric SRMs (e.g., for directed interactions) in which each individual contributes separately as "actor" and "partner" through correlated varying effects.

The Bhattacharyya coefficient (BC) of overlap is bounded on $[0, 1]$, and each estimate $\widehat{\mathrm{BC}}_{ij,t}$ for dyad $\{i, j\}$ in year $t$ has an associated 95% confidence interval derived from the autocorrelated kernel density estimation procedure [11]. To incorporate this estimation uncertainty into the model and to retain a tractable Gaussian likelihood, we logit-transformed each point estimate and its bounds, computed the standard error of the logit estimate from the transformed CI width as $\sigma_{ij,t} = (\mathrm{logit}(\widehat{\mathrm{BC}}^{U}_{ij,t}) - \mathrm{logit}(\widehat{\mathrm{BC}}^{L}_{ij,t}))/3.92$, and modelled the logit-transformed estimate with a measurement-error term [20]. The full specification is:

$$y_{ij,t} \sim \mathrm{Normal}(\mu_{ij,t},\ \sigma_{ij,t}) \tag{1}$$

$$\begin{aligned}\mu_{ij,t} = \bar{\alpha} + d_{\{ij\}} + \tfrac{1}{2}(m_i + m_j) + \beta_D\, D_{ij,t} + \beta_O\, (O_{i,t} + O_{j,t}) \\ + \beta_R\, R_{ij} + \beta_H\, H_{ij,t} + \beta_F\, F_t + \beta_E\, E_t\end{aligned} \tag{2}$$

where $y_{ij,t} = \mathrm{logit}(\widehat{\mathrm{BC}}_{ij,t})$ is the logit-transformed overlap estimate for dyad $\{i, j\}$ in year $t$, and $\sigma_{ij,t}$ is its known measurement standard error. The linear predictor $\mu_{ij,t}$ comprises:

- $\bar{\alpha}$: the overall intercept, representing the mean logit overlap across all dyads and years;
- $d_{\{ij\}}$: a dyad-level varying intercept capturing the unique pairwise deviation of dyad $\{i, j\}$ from the individual-level expectations;
- $m_i, m_j$: individual-level varying intercepts for the two mothers in dyad $\{i, j\}$, entered through the multi-membership structure $\frac{1}{2}(m_i + m_j)$, which assigns equal weight to both mothers and reflects the symmetric contribution of each individual to dyadic overlap;
- $D_{ij,t}$: absolute difference in offspring ages within the dyad in year $t$;
- $O_{i,t}, O_{j,t}$: the ages of mother $i$'s and mother $j$'s focal offspring in year $t$, entered as a single summed term $(O_{i,t} + O_{j,t})$ with a shared slope $\beta_O$, reflecting the symmetric contribution of each offspring's developmental stage to dyadic outcomes;
- $R_{ij}$: pedigree-based coefficient of relatedness between mothers $i$ and $j$;
- $H_{ij,t}$: mean home-range area of the two mothers in year $t$;
- $F_t, E_t$: mean fruit availability index (FAI) and Enhanced Vegetation Index (EVI) over the dyad-year sampling window.

The varying intercepts are modeled as independent draws from zero-centered normal distributions:

$$d_{\{ij\}} \sim \text{Normal}(0,\ \tau_d) \tag{3}$$
$$m_i \sim \text{Normal}(0,\ \tau_m) \tag{4}$$

where $\tau_d$ and $\tau_m$ are the among-dyad and among-mother standard deviations, respectively.

We placed weakly informative priors on all parameters [17]:

$$\bar{\alpha} \sim \text{Normal}(-0.55,\ 0.5) \tag{5}$$
$$\beta_D, \beta_O, \beta_R, \beta_H, \beta_F, \beta_E \sim \text{Normal}(0,\ 1) \tag{6}$$
$$\tau_d, \tau_m \sim \text{Exponential}(1) \tag{7}$$

The intercept prior was centred on the empirical mean of the logit-transformed BC estimates ($\approx -0.55$); and slope priors are weakly regularising on the standardised scale.

##### Symmetric SRM structure and its multiple membership implementation

Because our response is symmetric, we did not estimate the actor-partner correlation that is standard in asymmetric SRMs. Each individual contributes a single varying intercept through both "slots" of every dyad it participates in, and partial pooling across dyads that share an individual is achieved directly through the multi-membership structure rather than through a correlated bivariate random effect.

The specification above may be described as a multiple membership relations model [24], and is functionally equivalent to a symmetric (non-directed) SRM coded directly in a probabilistic programming language such as `Stan`; the two differ only in how the individual effects are constructed. A symmetric social relations model lacks a scalar or weight parameter that is multiplied by the individual-level varying effect as has been mathematically formalized (mostly in the supplementary material) in [23–26]. Coded directly, individuals are referenced by identity and the two members of a dyad enter as a sum of their effects, each with an implied coefficient of 1.

The multiple-membership model is functionally the same, but includes a weight value that the user can specify [27, 28], and is commonly used in various social network approaches (albeit with different weights or scalars)— see [29, 30]. `brms` instead uses a classic multiple membership model [27] which defines varying effects on data-frame columns, so each individual is split across the two member columns of a dyad; the multiple-membership term recombines these columns into a single per-individual effect and normalizes the membership weights to sum to one, so that each member enters with weight $\frac{1}{2}$ and the dyad's contribution is an average rather than a sum. This rescales the among-mother standard deviation $\tau_m$ by a constant factor but leaves predictions unchanged. In both formulations, the relationship effect is the dyad-level varying intercept $d_{\{ij\}}$. These model types are isomorphic and functionally make the same predictions– they just share different preferences for the use and/or magnitude of normalization constants.

##### Other dyadic and individual-level models

The other models in this study share components of the structure above and we describe them by their differences. The infant-infant play model retains the dyadic varying-effects structure (multi-membership term on mother identity and dyad-level varying intercept) and the same set of covariates (offspring age difference, summed offspring age, relatedness, FAI, and EVI), but is fitted to annual play-scan counts $N_{ij,t}$ using a hurdle Poisson likelihood (log link). The log of joint focal-scan observation effort is included as an offset, and a quadratic term in summed offspring age ($\beta_{O^2}\,(O_{i,t} + O_{j,t})^2$) is added to both the count and the hurdle components, reflecting the non-monotonic developmental trajectory of play previously documented in this population [31]. The hurdle component shares the same linear predictor as the count component but on the logit scale, with observation effort entered as a covariate rather than as an offset.

The models for daily path length and feeding time use individual-level structure: a single varying intercept per focal mother and focal offspring age, FAI, and EVI as covariates. The exposure variable was operationalized in two complementary specifications: a three-level predictor (alone with offspring; association without play; infant-infant play day) and a four-level adjacency predictor (no-play; day-before play; day-of play; day-after play). Daily path length is modelled with a Gamma likelihood (log link) and a measurement-error term derived from CTSD confidence intervals, and feeding effort with a beta-binomial likelihood (logit link) on counts of feeding scans out of total scans per follow day.

The Approach and Return models share a structure that differs from the dyadic and day-level models above. Rows are indexed at the (focal mother, partner, date) level, with a Gaussian likelihood on the change in distance to the reference core (partner's core for Approach, focal's own core for Return). With this dyad-indexed structure, each baseline focal-date contributes one row per potential play partner (the set of dyads in which the focal mother has been observed playing). This builds a counterfactual baseline distribution of the focal's daily travel for each (focal, partner) pair — what her travel looks like, relative to that specific partner's core, on days when no play is impending. The dyad random intercept accounts for non-independence among the multiple baseline observations sharing a focal-date by partially pooling within each dyad. The predictor of interest is a categorical phase variable contrasting baseline days with day-before and within-day pre-play windows (Approach) or with within-day post-play and day-after windows (Return). Covariates follow the DAG-based adjustment set described above (see DAG considerations for the directed-travel models), with random intercepts for focal mother and mother-dyad to account for repeated sampling within individuals and dyads. Although the dyad structure is reflected through the random intercept, the response measures only the focal mother's travel, with the partner serving as a geometric reference point rather than as a second actor (note: simultaneous nest-to-nest tracking of both mothers in a dyad is rarely available in the data).

### Supplementary Results

#### Offspring independently contribute to the age-difference effect on play

The play model presented in the main text uses total observation effort as the offset, estimating the total effect of offspring age difference on play frequency per unit of observation time. This estimand encompasses both mechanisms by which age difference might affect play: the indirect path through which mothers of similar-aged infants are in association more (creating opportunity for play), and the direct path through which similar-aged infants engage more once together in association. Note that association here plays the same role that home-range overlap plays in the dyadic DAG (Figure 4): both index the extent to which mothers create opportunity for their offspring to encounter one another. To isolate the second mechanism, we refit the same model structure with joint-association scans (where both mothers and their dependent infants were observed together) as the offset. The age-difference coefficient under this offset estimates the play rate per scan of opportunity for the infants to play, rather than per scan of observation time. The effect remained substantially negative under both specifications ($\beta_{\text{association}}$ = -0.47 [-0.70, -0.25]; $PP > 0$ = 0; $\beta_{\text{observation}}$ = -0.33 [-0.57, -0.09]; $PP > 0$ = 0.011; Figure 9a). The persistence and stability of the age-difference effect across these specifications indicates that offspring themselves contribute substantively to the pattern: similar-aged infants engage in more play once their mothers have brought them into proximity.

#### Mothers independently contribute to the age-difference effect on play

The previous analysis isolates the offspring contribution but does not isolate the maternal contribution to the age-difference effect on play. Two further lines of evidence point to a substantial maternal contribution operating through space use. First, the main-text home-range overlap result shows that mother-mother spatial overlap (Bhattacharyya coefficient) is credibly higher for dyads with similar-aged offspring, indicating that mothers structure their intensive space use to bring similar-aged offspring into closer spatial proximity. Second, we fitted a hurdle negative-binomial symmetric SRM with mother-mother association scans as the outcome, joint observation effort as the offset, and the same environmental and demographic covariates as the other dyadic models. Mother relatedness had a credibly positive effect on association rate ($\beta$ = 0.43 [0.12, 0.76]; $PP > 0$ = 0.984; Figure 9b), replicating the kin-biased association pattern reported by [3]. Offspring age difference showed a weakly negative effect ($\beta$ = -0.18 [-0.42, 0.08], $PP > 0$ = 0.126) independent of mother relatedness. Third, we show in the main text that mothers oriented travel toward the partner's core on the day before play. Together, these findings suggest that mothers play an active role in shaping the social environment of their offspring.

#### Disentangling the costs of play from maternal association

To determine whether energetic costs reflected play itself or association with another mother, we refit the daily path length and feeding-time models with a four-level predictor distinguishing days with no association, days with non-mother association only, days with mother-mother association but no play, and play days (Figure 10). Daily travel on mother-mother association days without play was comparable to days with non-mother associates ($\beta$ = 0.03 [-0.05, 0.10]; $PP > 0 = 0.708$), but mothers travelled credibly farther on play days than on either mother-mother association days ($\beta$ = 0.14 [0.04, 0.22]; $PP > 0 = 0.992$) or other-association days ($\beta$ = 0.16 [0.09, 0.24]; $PP > 0 = 1.000$), consistent with mothers actively travelling farther to arrange infant play encounters. Feeding costs were more closely tied to maternal association. Mothers fed less on mother-mother association days than on other-association days ($\beta$ = -0.08 [-0.16, -0.01]; $PP > 0 =$ 0.046), and feeding declined further on play days, although the contrast between play and mother-mother association days remained uncertain ($\beta$ = -0.04 [-0.15, 0.05]; $PP > 0 = 0.248$). Together, these results suggest that elevated daily travel is specific to play, whereas reduced feeding is more closely tied to association with another mother, the social context that makes infant play possible. The weak additional reduction on play days may reflect time or attention diverted toward monitoring active play bouts, although this interpretation remains tentative.

#### Sensitivity to mixed association status in play-adjacency models

The play-adjacency models in the main text use a “no play” reference category that includes both fully-alone days and days when the focal mother associated with a non-offspring conspecific without infant play. Because non-offspring association can itself elevate travel and reduce feeding, we conducted a sensitivity analysis in which the no-play reference and the day-before/day-after categories were restricted to no-association days. Day-of-play observations are association days by definition and were unaffected. Within the original sample, the proportion of association days was 25.9% for the no-play reference, 27.7% for day-before, and 41.2% for day-after; the day-before category was therefore comparable to the reference, while day-after contained somewhat more association days. The filtered model (Tables 1 and 2) yielded effects in the same direction and of comparable magnitude to the original, including for day-after, supporting the interpretation that the adjacency gradient is not an artifact of association mixed into the “non-play day” categories. We retain the original specification in the main text because it preserves the full sample of adjacency days and is consistent with the temporal scale of the question (whether the days surrounding play differ from ordinary days), while the filtered model serves as a robustness check.

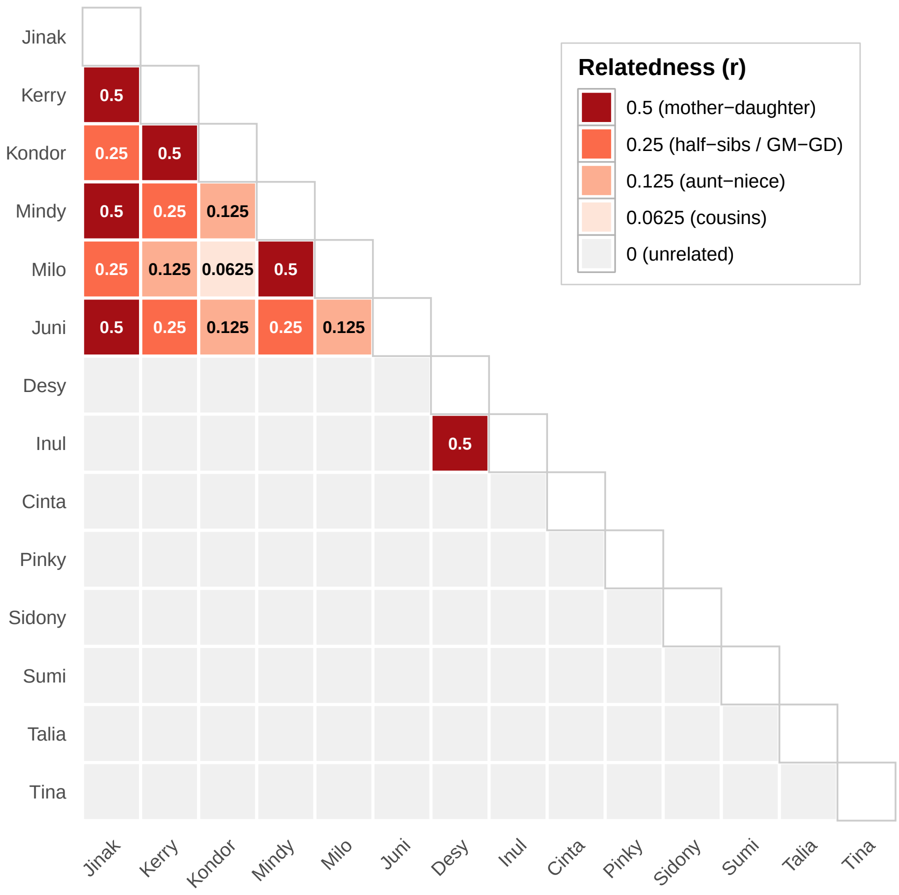


Figure 1: **Maternal pedigree relatedness ($r$) among the 14 mothers included in the Tuanan dyadic analyses.** Each cell shows the coefficient of relatedness for one female-female dyad, derived from long-term maternal pedigrees of the Tuanan Orangutan Research Project. Cell color encodes relatedness category: $r = 0.5$ for mother-daughter; $r = 0.25$ for maternal half-sisters and grandmother-granddaughter pairs (both share, on average, a quarter of their genome through the maternal line); $r = 0.125$ for aunt-niece; $r = 0.0625$ for maternal cousins; and $r = 0$ for pairs with no documented common maternal ancestor. Cells along the diagonal are left blank because relatedness is not defined for an individual to herself. Females are ordered to keep matrilineally related individuals adjacent: the upper-left block comprises Jinak's matriline (her daughters Kerry, Mindy, Juni; her granddaughters Kondor and Milo), followed by the Desy-Inul mother-daughter pair, with the remaining six females belonging to separate matrilines. The matrix shows all 91 possible pairs among these 14 mothers; 71 pairs contributed at least one dyad-year to the analysis (282 dyad-years total), with the remainder absent because the two mothers were never simultaneously followed in a year when both had a dependent offspring.

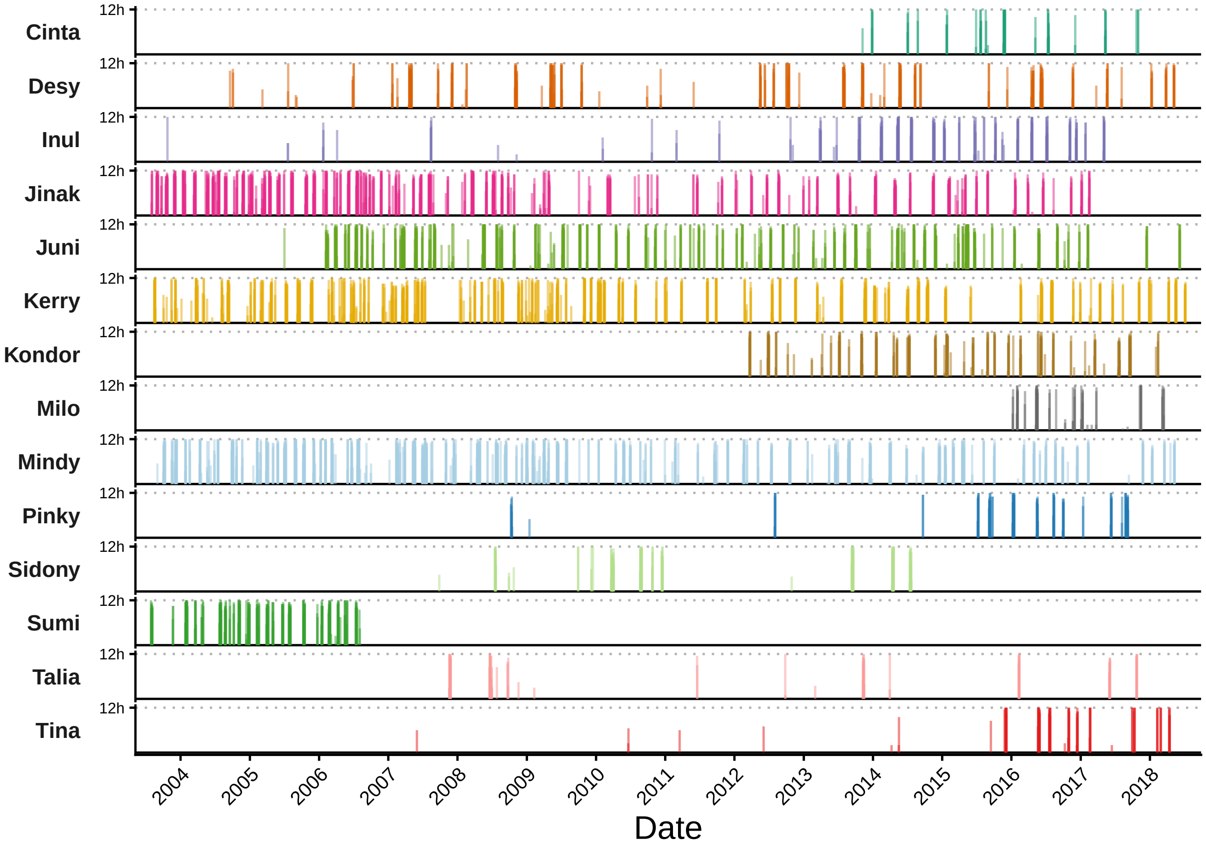


Figure 2: **GPS sampling coverage for focal orangutan mothers, 2003-2018.** Each row shows daily tracking coverage for one female ($n = 14$) included in downstream analyses of daily path length and/or dyadic home-range overlap. Vertical bars indicate the proportion of a full follow day (12 h, equivalent to 24 fixes at 30-min sampling) covered on each date. Dotted horizontal lines mark full-day coverage. Coverage gaps reflect periods when females were not the focal subject of nest-to-nest follows.

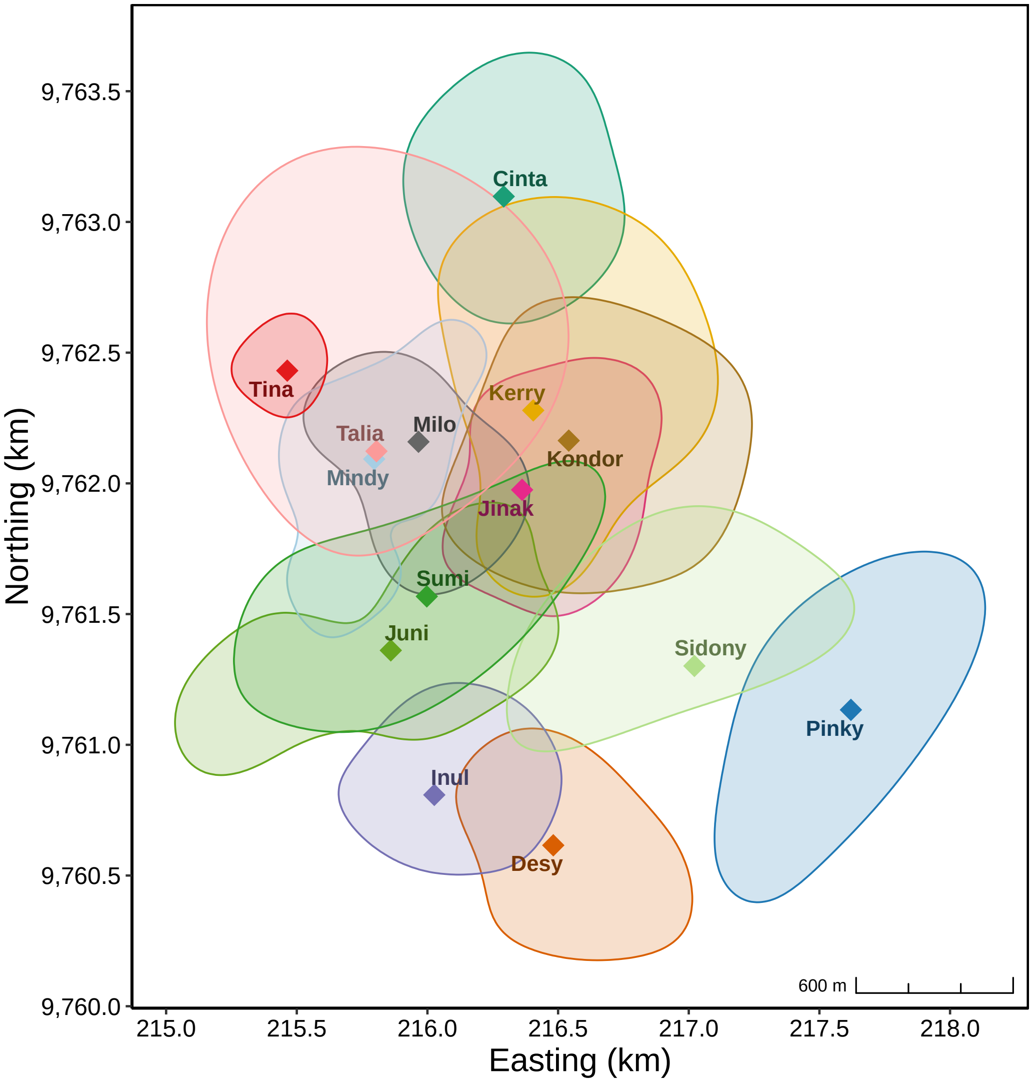


Figure 3: **Core areas and mean locations of focal orangutan mothers, 2003-2018.** Polygons show each female's 50% core area, estimated from their full study-period autocorrelated kernel density estimate (AKDE). Diamonds mark each female's all-time mean location.

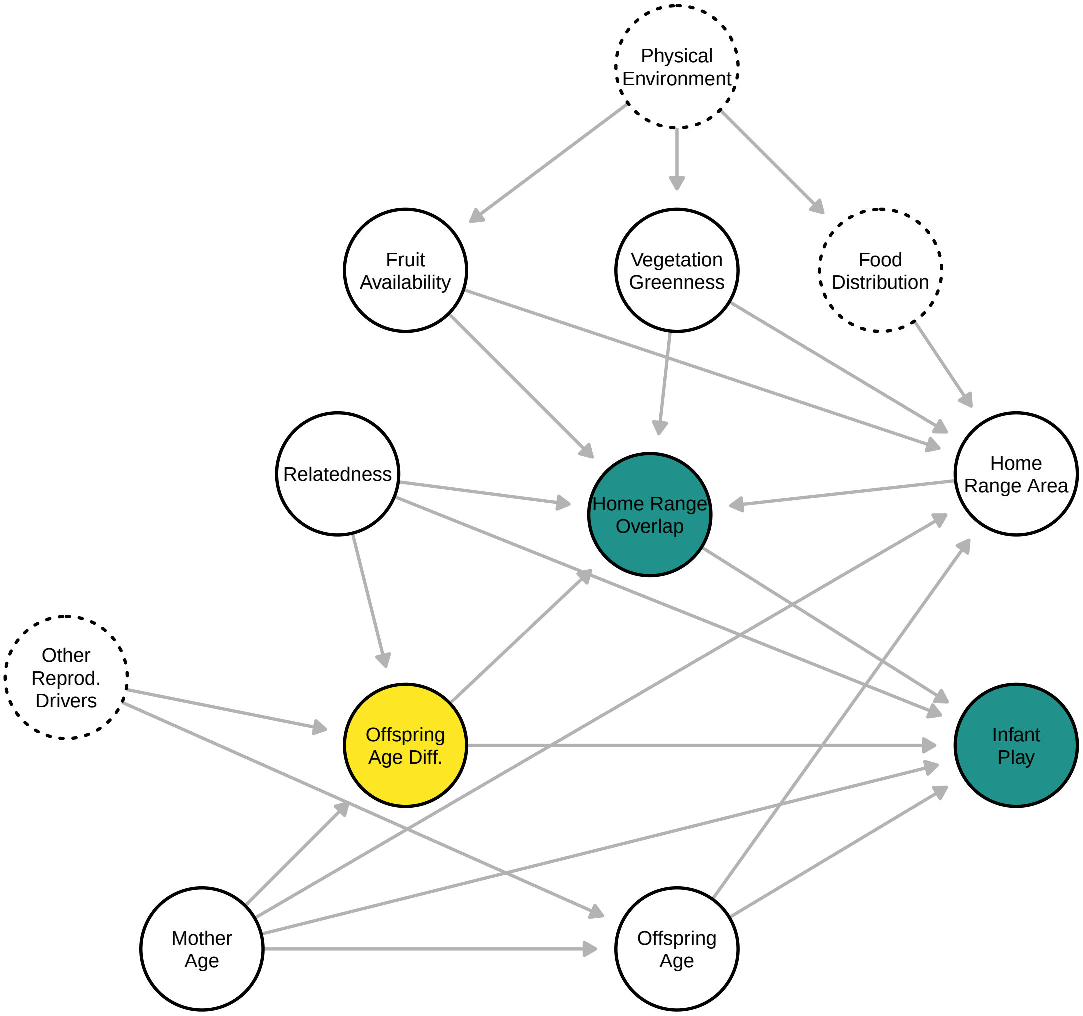


Figure 4: **Directed acyclic graph for the dyadic models** (home-range overlap, intersection area, and infant-infant play). Solid circles denote observed variables; dashed circles denote latent variables. Yellow: exposure (OFF_AGE_DIFF). Teal: outcomes (OVER, PLAY); the same DAG applies to intersection area. See SI Materials and Methods for variable definitions, arrow justifications, and adjustment sets.

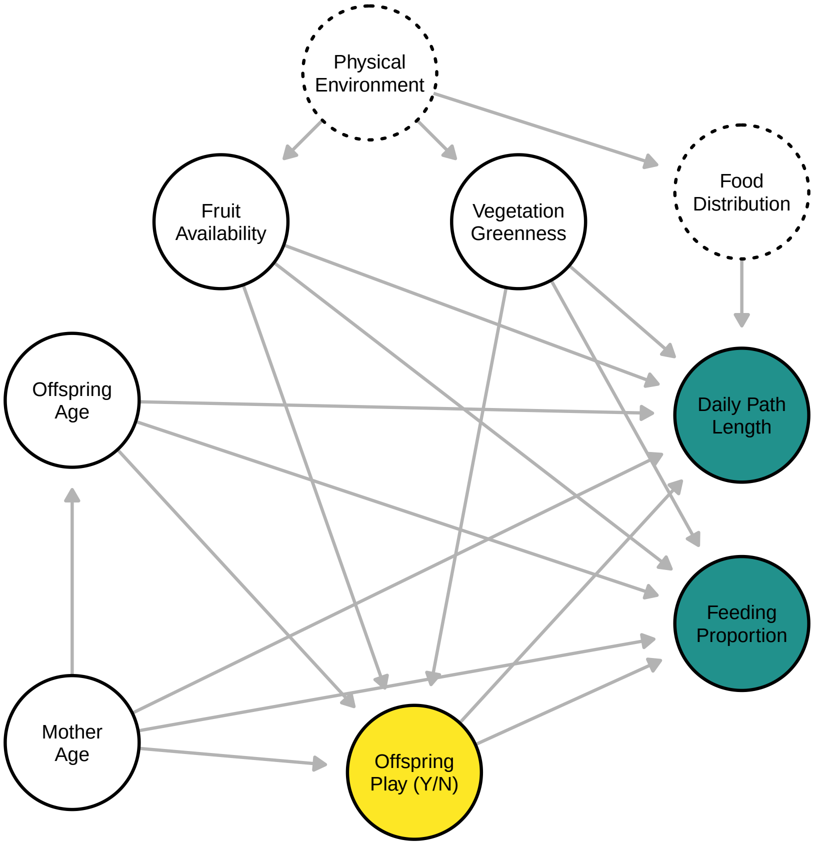


Figure 5: **Directed acyclic graph for the day-level models** (daily path length and feeding proportion). Solid circles denote observed variables; dashed circles denote latent variables. Yellow: exposure (PLAY). Teal: outcomes (DPL, FEED). See SI Materials and Methods for variable definitions, arrow justifications, and adjustment sets.

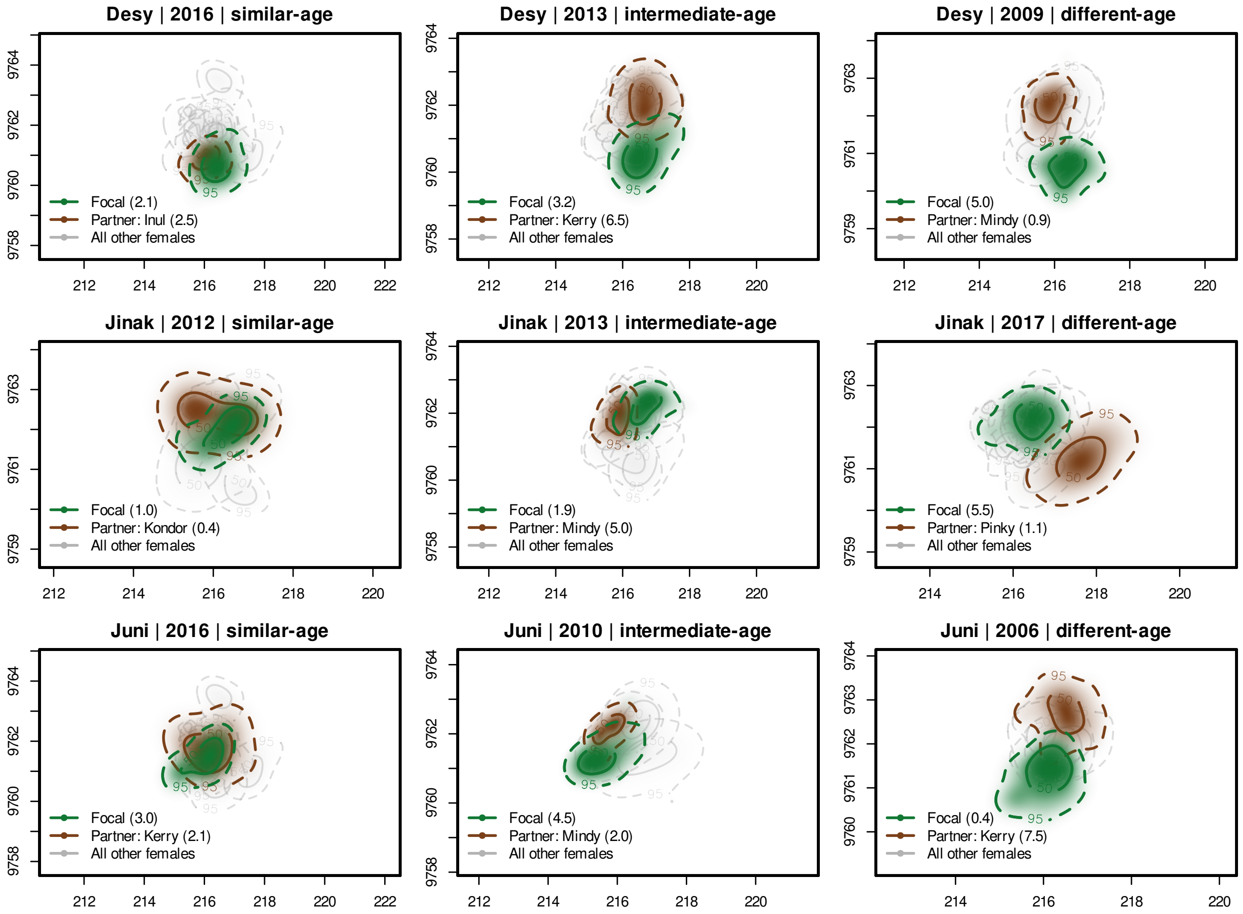


Figure 6: **Examples of annual home-range overlap varying with offspring age similarity.** Selected dyad-years illustrating the spatial pattern underlying the results in Figure 1. Each row shows a focal mother (green) paired with three different neighbors (brown) in separate dyad-years representing similar (left column), intermediate (middle column), and large (right column) offspring age differences. Shaded regions show 95% and 50% autocorrelated kernel density estimate (AKDE) contours, with underlying color gradients indicating the probability of space use. Light grey contours show home ranges of all other mothers in the study population for that year. Offspring ages for each dyad are shown in the panel legends.

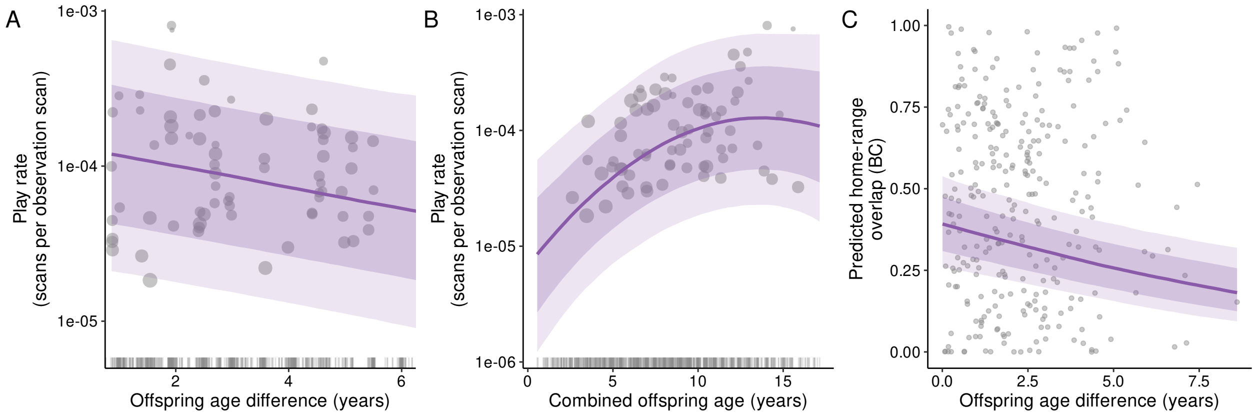


Figure 7: **Model predictions for the effect of offspring age on infant-infant play and on maternal home-range overlap.** (a) Predicted play rate (play scans per observation scan) as a function of the difference in offspring age within a infant-infant dyad. (b) Predicted play rate as a function of combined offspring age (sum of the two infants' ages), capturing the quadratic age effect. (c) Predicted home-range overlap (Bhattacharyya coefficient) as a function of offspring age difference. In all panels the solid line is the posterior median trend and the shaded bands show the 66% and 89% posterior credible intervals, with all other predictors held at their mean. Grey points are the raw dyad-year data; point size in (a) and (b) is scaled to observation effort. Play rate is shown on a logarithmic axis; dyad-years with zero observed play (the majority) cannot be displayed on this axis and are instead shown as a rug along the lower margin of (a) and (b).

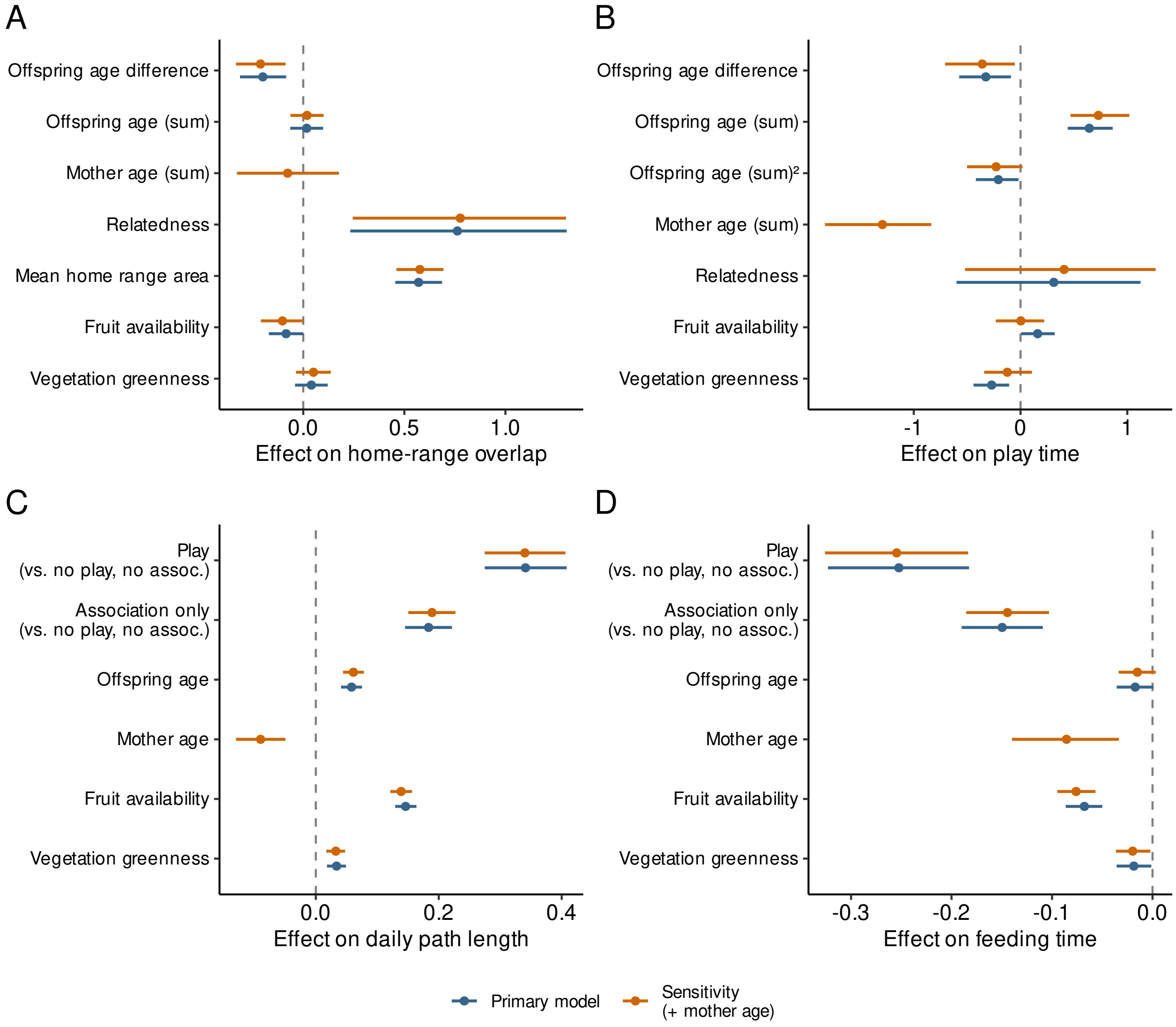


Figure 8: **Sensitivity of dyadic- and individual-level effects to the inclusion of maternal age.** 89% posterior point intervals for fixed-effect coefficients in the primary models (blue) and sensitivity models (orange) for (a) home-range overlap, (b) infant–infant play time, (c) daily path length, and (d) feeding time. Sensitivity models include summed maternal age (a, b) or maternal age (c, d) as an additional covariate, entered via a measurement-error specification [20] that propagates encoding uncertainty in birth-year estimates (most mothers had only upper-bound birth years; see Methods). For the play model (b), coefficients shown are from the count component of the hurdle, describing effects on the rate of play conditional on any play occurring.

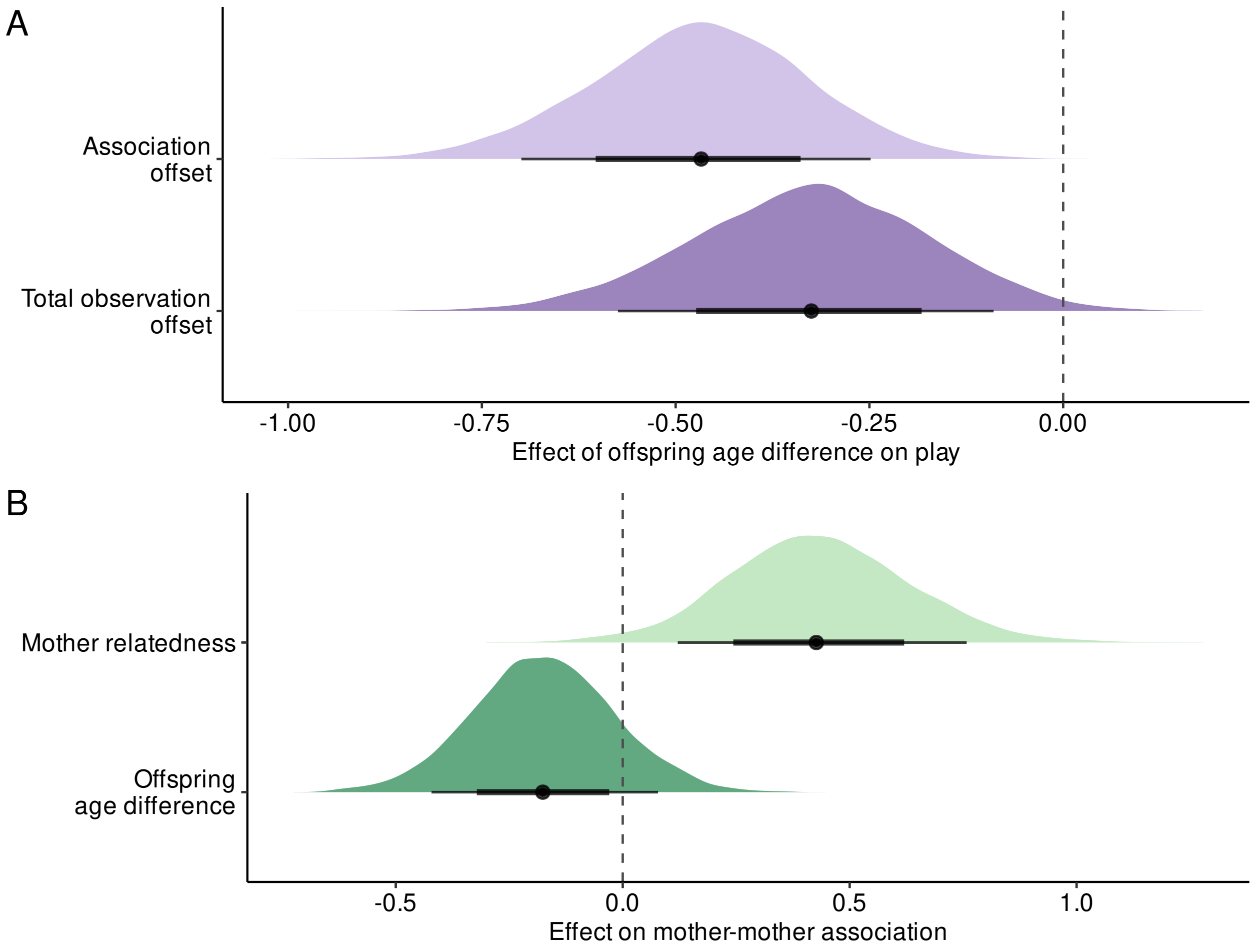


Figure 9: **Decomposition of the age-difference effect on play.** *(a)* Posterior estimates of the offspring age-difference coefficient in the play-scan model under two offset specifications. Using total observation scans as the offset estimates the total effect of age difference on play, whereas using association scans conditions on the joint presence of the two mothers and their dependent offspring. *(b)* Posterior estimates of the mother relatedness and offspring age-difference coefficients in the mother-mother association model. Points show posterior medians, and thick and thin lines show 66% and 89% credible intervals, respectively.

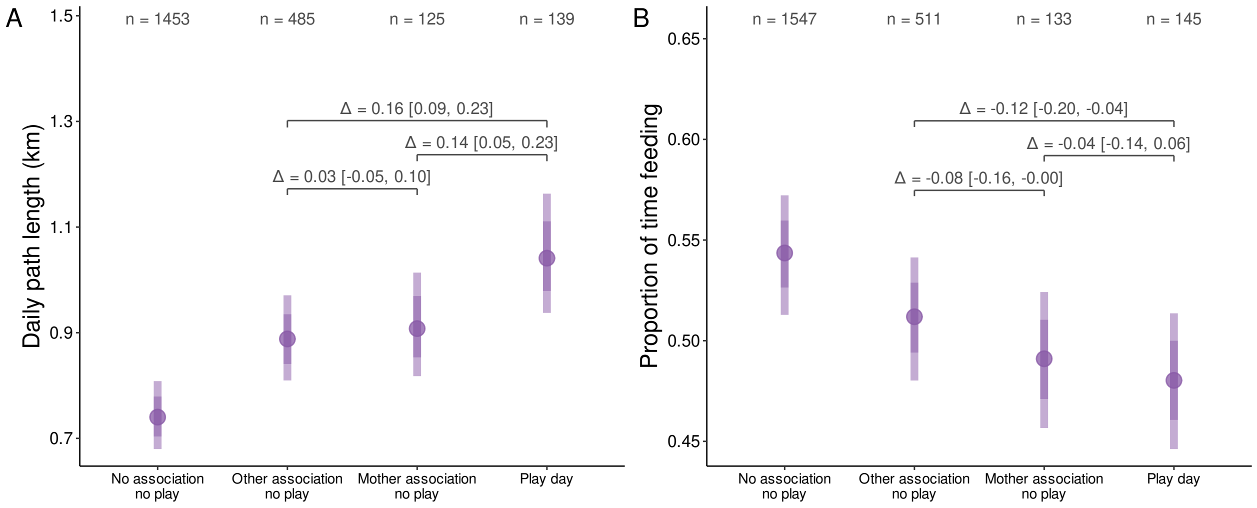


Figure 10: **Disentangling the costs of play from maternal association.** Posterior predictions from Bayesian GLMMs of daily path length (*a*) and proportion of time feeding (*b*), comparing four categories: days when the focal mother was alone with her offspring (no association, no play; reference), days when she associated only with non-mother conspecifics without infant play (other association, no play), days when she associated with another mother without infant play (mother association, no play), and days when infant-infant play occurred (play day; which by definition involves mother-mother association). Posterior predicted values across categories are shown with covariates held at their mean values; sample sizes ($n$) are shown for each category. Points and dark/light lines show posterior medians with 66% and 89% credible intervals. Brackets and labelled values within each panel indicate the posterior median and 89% credible interval of the contrast between play days and the two association-only categories.

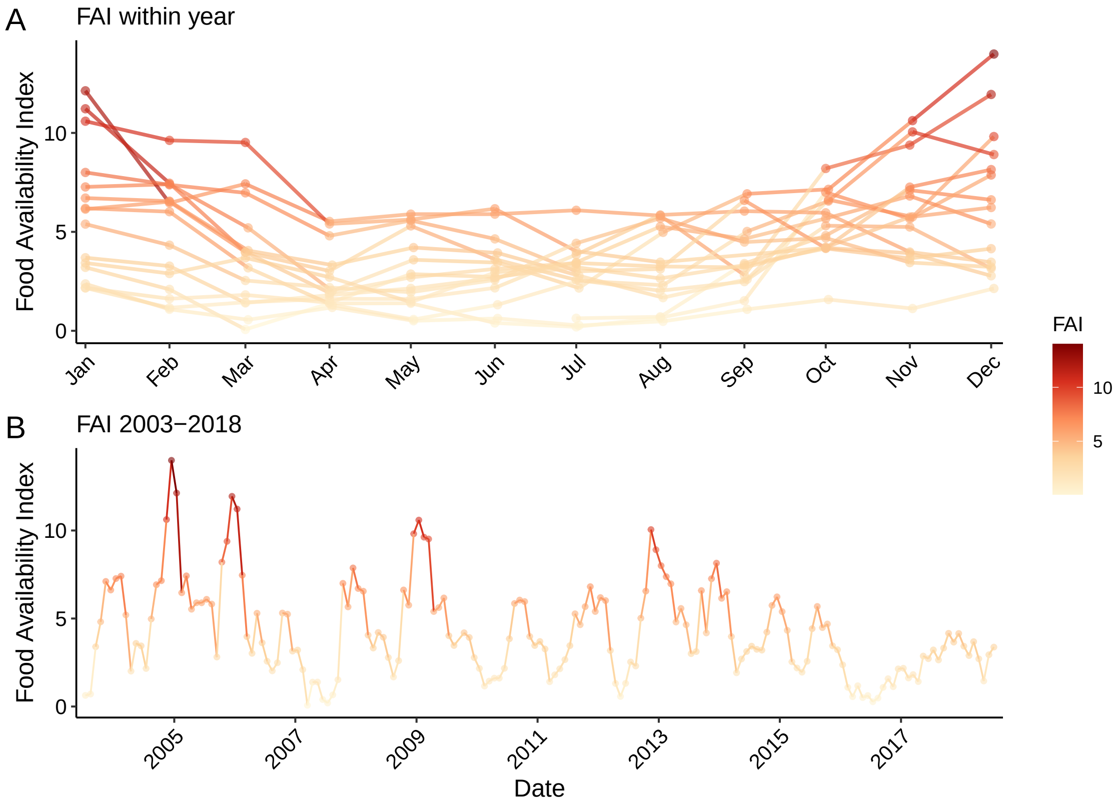


Figure 11: **Fruit Availability Index (FAI) at Tuanan, 2003-2018.** FAI was scored monthly from phenological monitoring of marked trees along established transects at the field site. (*a*) Within-year seasonal variation: each line represents one calendar year. (*b*) Full monthly time series, 2003-2018. Colour reflects FAI value across both panels.

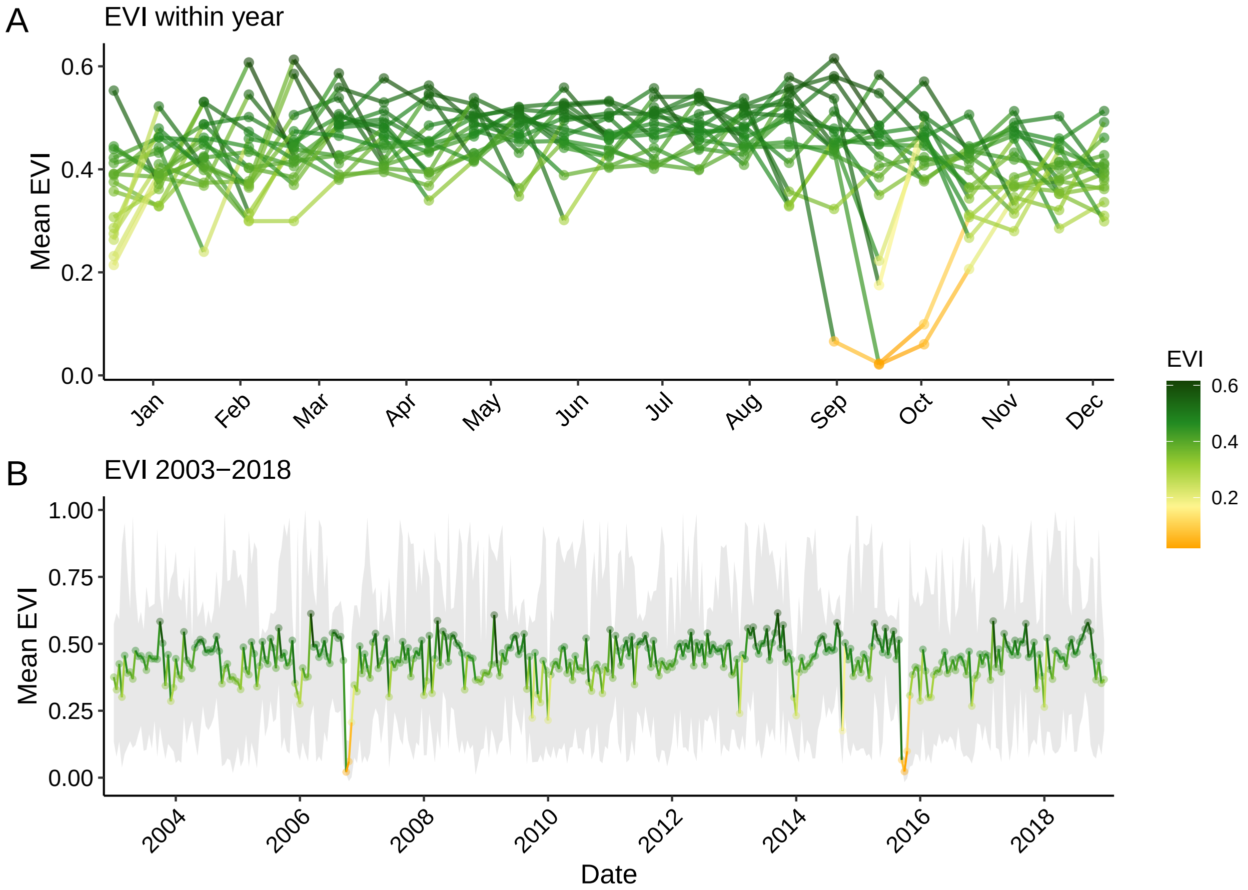


Figure 12: **Enhanced Vegetation Index (EVI) at Tuanan, 2003-2018.** ($a$) Within-year seasonal variation: each line represents one calendar year, with points showing 16-day composite values. ($b$) Full time series, 2003-2018, showing the spatial mean across pixels (line and points) and the spatial range across the window (grey ribbon). Colour reflects EVI value across both panels.

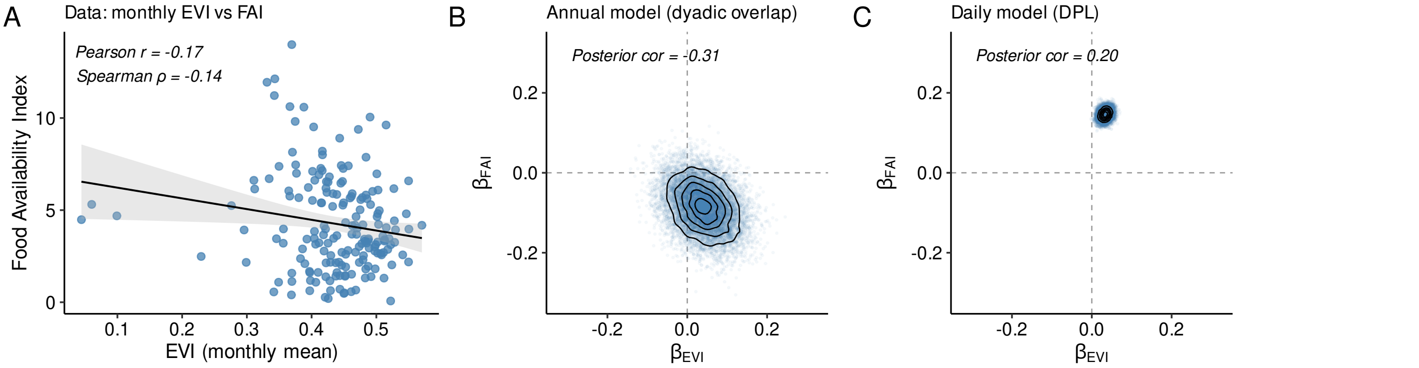


Figure 13: **EVI and FAI capture distinct sources of environmental variation.** ($a$) Monthly mean EVI plotted against monthly habitat-wide FAI (2003-2018). The black line shows a linear regression with 95% confidence band; Pearson and Spearman correlations are shown for descriptive purposes only. ($b$, $c$) Joint posterior distributions of the standardized FAI and EVI regression coefficients in the annual dyadic-overlap model ($b$) and the daily path-length model ($c$). Blue points are individual posterior draws; black contours show 2D density. Dashed lines mark zero on each axis. The posterior correlation between the two coefficients is reported in each panel; roughly circular posteriors indicate that each effect was independently identifiable in the model.

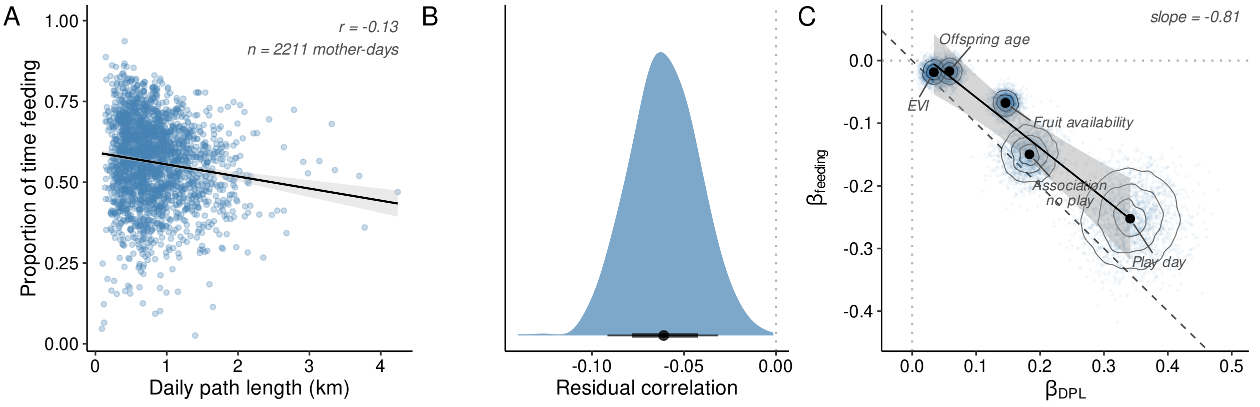


Figure 14: **Daily path length and feeding rate are coupled but capture partly independent aspects of mothers' activity budgets.** *(a)* Daily proportion of time spent feeding plotted against daily path length, with one point per mother-day ($n = 2216$). The black line and grey ribbon show a linear regression with a 95% confidence band. The two outcomes are weakly negatively correlated ($r = -0.14$). *(b)* Posterior distribution of the residual correlation between the two model fits, computed by correlating posterior residuals across mother-days within each posterior draw (median $r = -0.06$, 89% HDI $[-0.09, -0.04]$). *(c)* For each predictor that appears in both models, the joint posterior of its DPL coefficient ($\beta_{DPL}$, x-axis) against its feeding coefficient ($\beta_{feeding}$, y-axis), shown as 2D density contours with black points at the posterior medians. The dashed grey line marks $y = -x$, where the points would fall if the two models carried identical information with opposite signs. The solid blackline and grey ribbon show a linear regression with a 95% confidence band through the five coefficient medians (slope $= -0.81$). All five coefficients sit above the $y = -x$ reference, meaning that for every predictor, daily path length shifts more than feeding rate does.

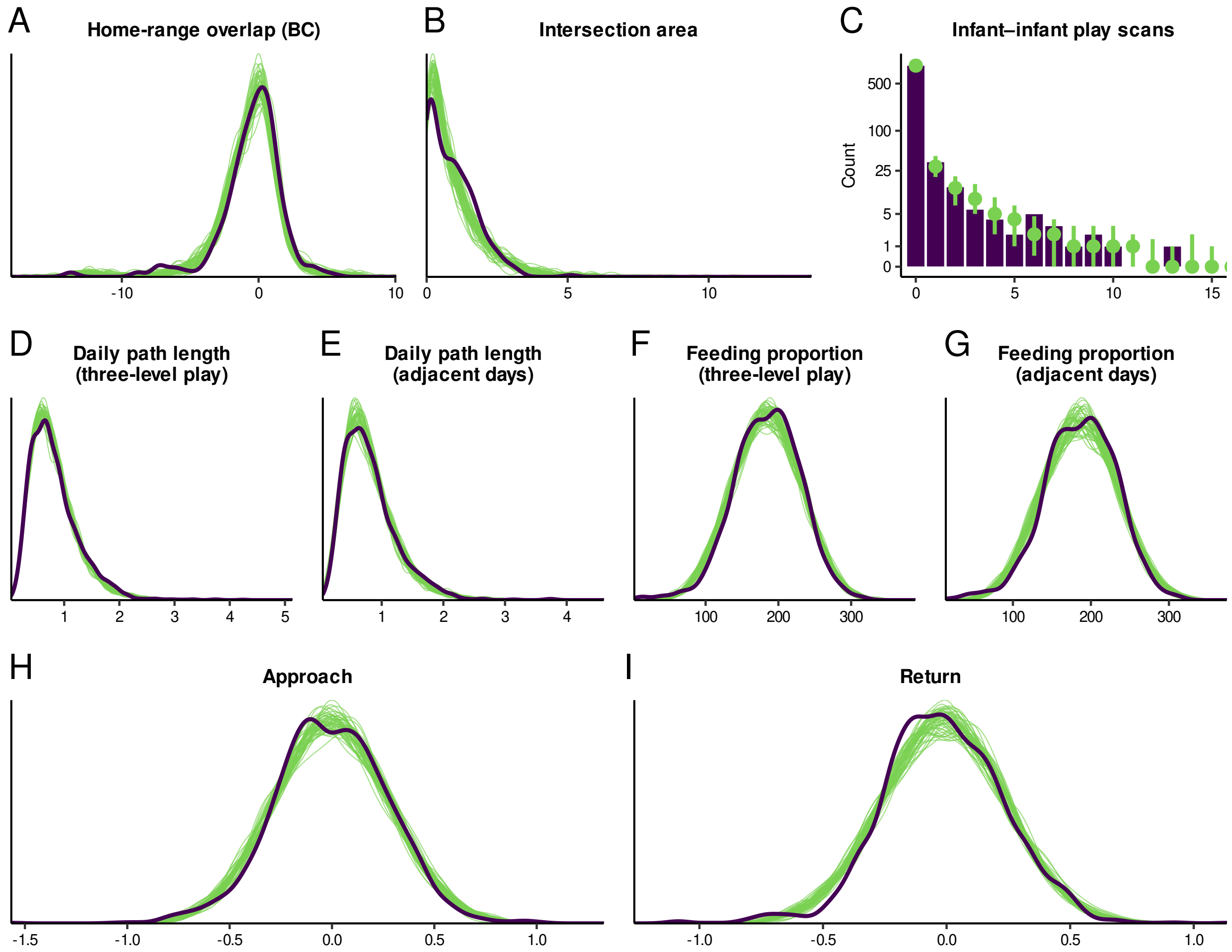


Figure 15: **Posterior predictive checks for all primary Bayesian models.** Panels *a*, *b*, and *d*-*i* show the observed outcome distribution (dark purple line) overlaid with 50 draws from the posterior predictive distribution (green lines). Panel *c* shows a posterior-predictive bar plot with log-transformed y-axis rather than a density overlay because the play-scan outcome is a discrete count with substantial zero-inflation; observed counts at each value are shown as dark purple bars with posterior-predictive medians and intervals overlaid in green.

#### Sensitivity summaries for the play-adjacency models

The following tables compare the original and filtered specifications of the play-adjacency models (see SI Results for details).

Table 1: Sensitivity comparison of the daily path length adjacency model. Original (orig.) includes association days; filtered (filt.) restricts day_before and day_after to no-association days. Columns: posterior median with 89% HDI, $P(\beta > 0)$, sample size, and % association days in the original sample.

| Category | $\beta$ (orig.) | $P(\beta > 0)$ orig. | n (orig.) | % assoc. | $\beta$ (filt.) | $P(\beta > 0)$ filt. | n (filt.) |
|---|---|---|---|---|---|---|---|
| Day before play | 0.09 [-0.02, 0.20] | 0.90 | 46 | 28.3 | 0.09 [-0.04, 0.22] | 0.86 | 33 |
| Day of play | 0.34 [0.27, 0.41] | 1.00 | 139 | 100.0 | 0.40 [0.33, 0.47] | 1.00 | 139 |
| Day after play | 0.14 [0.05, 0.23] | 0.99 | 70 | 42.9 | 0.23 [0.12, 0.35] | 1.00 | 40 |

Table 2: Sensitivity comparison of the feeding-proportion adjacency model. Columns as in Table 1.

| Category | $\beta$ (orig.) | $P(\beta > 0)$ orig. | n (orig.) | % assoc. | $\beta$ (filt.) | $P(\beta > 0)$ filt. | n (filt.) |
|---|---|---|---|---|---|---|---|
| Day before play | 0.03 [-0.09, 0.15] | 0.68 | 49 | 26.5 | 0.04 [-0.09, 0.18] | 0.68 | 36 |
| Day of play | -0.23 [-0.31, -0.16] | 0.00 | 145 | 100.0 | -0.28 [-0.35, -0.20] | 0.00 | 145 |
| Day after play | -0.04 [-0.13, 0.07] | 0.28 | 75 | 42.7 | -0.00 [-0.13, 0.12] | 0.49 | 43 |

#### Posterior summaries for all models

The following tables provide posterior medians, 89% highest density intervals (HDIs), $\hat{R}$, and bulk and tail effective sample sizes (ESS) for all parameters in each of the nine primary Bayesian models reported in this study.

Table 3: Posterior summaries for the home-range overlap (BC) model. Median, 89% highest density interval (HDI), $\hat{R}$, and bulk and tail effective sample sizes (ESS) for each parameter.

| Parameter | Median | HDI lo | HDI hi | Rhat | ESS bulk | ESS tail |
|---|---|---|---|---|---|---|
| $\beta$ Vegetation greenness (EVI) | 0.04 | -0.04 | 0.12 | 1.001 | 17880 | 10033 |
| $\beta$ Fruit availability | -0.08 | -0.17 | 0.00 | 1.000 | 17488 | 9424 |
| Intercept (count) | -0.72 | -1.28 | -0.14 | 1.000 | 12551 | 8689 |
| $\beta$ Mean home-range area | 0.57 | 0.46 | 0.69 | 1.000 | 18045 | 8865 |
| $\beta$ Offspring age difference | -0.20 | -0.32 | -0.09 | 1.000 | 17372 | 9490 |
| $\beta$ Offspring age (sum) | 0.02 | -0.06 | 0.10 | 1.000 | 16548 | 8837 |
| $\beta$ Relatedness | 0.76 | 0.24 | 1.31 | 1.000 | 6211 | 7283 |
| SD (dyad) | 1.87 | 1.57 | 2.22 | 1.000 | 3508 | 5486 |
| SD (mother) | 1.72 | 0.82 | 2.71 | 1.000 | 2665 | 2291 |

Table 4: Posterior summaries for the home-range intersection-area model (hurdle Gamma).

| Parameter | Median | HDI lo | HDI hi | Rhat | ESS bulk | ESS tail |
|---|---|---|---|---|---|---|
| $\beta$ Vegetation greenness (EVI) | 0.03 | -0.05 | 0.10 | 1.000 | 12872 | 9349 |
| $\beta$ Fruit availability | -0.03 | -0.11 | 0.06 | 1.000 | 11896 | 9201 |
| $\beta_{hu}$ Vegetation greenness (EVI) | -0.36 | -0.77 | 0.03 | 1.000 | 14752 | 8717 |
| $\beta_{hu}$ Fruit availability | -0.48 | -0.93 | -0.04 | 1.000 | 15796 | 8693 |
| Intercept (hurdle) | -4.33 | -6.18 | -2.85 | 1.000 | 8458 | 4631 |
| $\beta_{hu}$ Mean home-range area | -1.00 | -1.43 | -0.56 | 1.001 | 18902 | 9251 |
| $\beta_{hu}$ Offspring age difference | 0.04 | -0.43 | 0.52 | 1.000 | 19059 | 9572 |
| $\beta_{hu}$ Offspring age (sum) | 0.32 | -0.05 | 0.72 | 1.000 | 17929 | 9172 |
| $\beta_{hu}$ Relatedness | -3.33 | -5.97 | -1.27 | 1.000 | 8675 | 4792 |
| Intercept (count) | -0.31 | -0.64 | 0.01 | 1.000 | 6365 | 6850 |
| $\beta$ Mean home-range area | 0.42 | 0.34 | 0.50 | 1.000 | 15073 | 9486 |
| $\beta$ Offspring age difference | -0.06 | -0.15 | 0.03 | 1.000 | 15052 | 8886 |
| $\beta$ Offspring age (sum) | 0.00 | -0.08 | 0.07 | 1.000 | 16212 | 8690 |
| $\beta$ Relatedness | 0.48 | 0.31 | 0.65 | 1.000 | 6897 | 8177 |
| Intercept (shape) | 1.05 | 0.89 | 1.21 | 1.000 | 7296 | 8377 |
| $\beta_{shape}$ Mean home-range area | 0.56 | 0.39 | 0.74 | 1.000 | 12011 | 9201 |
| SD (dyad) | 0.52 | 0.35 | 0.69 | 1.000 | 2417 | 5001 |
| SD (mother) | 0.64 | 0.11 | 1.08 | 1.001 | 1761 | 2070 |

Table 5: Posterior summaries for the infant–infant play-scan model (hurdle Poisson). $\beta$ rows refer to the count component; $\beta_{hu}$ rows refer to the hurdle (zero) component.

| Parameter | Median | HDI lo | HDI hi | Rhat | ESS bulk | ESS tail |
|---|---|---|---|---|---|---|
| $\beta$ Offspring age difference | -0.32 | -0.57 | -0.08 | 1.000 | 11292 | 9506 |
| $\beta$ Offspring age (sum) | 0.64 | 0.44 | 0.86 | 1.000 | 9130 | 8336 |
| $\beta$ Vegetation greenness (EVI) | -0.27 | -0.44 | -0.10 | 1.000 | 11260 | 8835 |
| $\beta$ Fruit availability | 0.16 | 0.01 | 0.32 | 1.000 | 11872 | 9526 |
| $\beta_{hu}$ Offspring age difference | 0.28 | -0.03 | 0.58 | 1.000 | 10621 | 9037 |
| $\beta_{hu}$ Offspring age (sum) | -0.90 | -1.21 | -0.60 | 1.000 | 9489 | 8665 |
| $\beta_{hu}$ Vegetation greenness (EVI) | 0.00 | -0.29 | 0.29 | 1.000 | 11498 | 9271 |
| $\beta_{hu}$ Fruit availability | -0.32 | -0.61 | -0.02 | 1.000 | 11504 | 9030 |
| $\beta_{hu}$ Offspring age $(\text{sum})^2$ | 0.36 | 0.08 | 0.66 | 1.000 | 10133 | 8902 |
| Intercept (hurdle) | 19.55 | 15.35 | 23.95 | 1.000 | 7516 | 8348 |
| $\beta_{hu}$ log(observation effort) | -1.73 | -2.17 | -1.30 | 1.000 | 7852 | 8409 |
| $\beta_{hu}$ Relatedness | -1.14 | -1.34 | -0.95 | 1.000 | 10369 | 8129 |
| $\beta$ Offspring age $(\text{sum})^2$ | -0.21 | -0.41 | -0.01 | 1.000 | 10281 | 8754 |
| Intercept (count) | -1.38 | -3.10 | 0.29 | 1.000 | 10445 | 8343 |
| $\beta$ Relatedness | 0.31 | -0.55 | 1.18 | 1.000 | 3603 | 3159 |
| SD (dyad) | 0.48 | 0.00 | 1.36 | 1.002 | 1687 | 2089 |
| SD (dyad) | 0.48 | 1.48 | 1.52 | 1.002 | 1687 | 2089 |
| SD (mother) | 7.80 | 5.21 | 10.74 | 1.000 | 3325 | 4944 |

Table 6: Posterior summaries for the daily path length model (three-level association/play encoding).

| Parameter | Median | HDI lo | HDI hi | Rhat | ESS bulk | ESS tail |
|---|---|---|---|---|---|---|
| $\beta$ Vegetation greenness (EVI) | 0.03 | 0.02 | 0.05 | 1.000 | 22701 | 9286 |
| $\beta$ Fruit availability | 0.15 | 0.13 | 0.16 | 1.000 | 21883 | 10187 |
| Intercept (count) | -0.31 | -0.40 | -0.23 | 1.001 | 3248 | 5445 |
| $\beta$ Offspring age | 0.06 | 0.04 | 0.07 | 1.000 | 21621 | 9565 |
| $\beta$ Association, no play | 0.18 | 0.14 | 0.22 | 1.000 | 22655 | 9765 |
| $\beta$ Play day | 0.34 | 0.28 | 0.41 | 1.000 | 23479 | 9304 |
| SD (mother) | 0.18 | 0.12 | 0.25 | 1.000 | 4257 | 6972 |
| Shape | 5.01 | 4.77 | 5.26 | 1.000 | 23020 | 9036 |

Table 7: Posterior summaries for the daily path length model (four-level adjacent-day encoding).

| Parameter | Median | HDI lo | HDI hi | Rhat | ESS bulk | ESS tail |
|---|---|---|---|---|---|---|
| $\beta$ Vegetation greenness (EVI) | 0.05 | 0.03 | 0.08 | 1 | 15632 | 8847 |
| $\beta$ Fruit availability | 0.16 | 0.14 | 0.18 | 1 | 15216 | 9367 |
| Intercept (count) | -0.33 | -0.43 | -0.23 | 1 | 2519 | 4059 |
| $\beta$ Offspring age | 0.05 | 0.02 | 0.07 | 1 | 14834 | 9484 |
| $\beta$ Day after play | 0.14 | 0.05 | 0.23 | 1 | 17108 | 9410 |
| $\beta$ Day before play | 0.09 | -0.02 | 0.21 | 1 | 16699 | 8846 |
| $\beta$ Day of play | 0.34 | 0.27 | 0.41 | 1 | 14613 | 8672 |
| SD (mother) | 0.21 | 0.13 | 0.29 | 1 | 4133 | 6684 |
| Shape | 5.01 | 4.67 | 5.36 | 1 | 16453 | 9432 |

Table 8: Posterior summaries for the feeding-proportion model (three-level association/play encoding).

| Parameter | Median | HDI lo | HDI hi | Rhat | ESS bulk | ESS tail |
|---|---|---|---|---|---|---|
| $\beta$ Vegetation greenness (EVI) | -0.02 | -0.04 | 0.00 | 1 | 11848 | 8540 |
| $\beta$ Fruit availability | -0.07 | -0.09 | -0.05 | 1 | 11855 | 8619 |
| Intercept (count) | 0.16 | 0.05 | 0.27 | 1 | 2123 | 3379 |
| $\beta$ Offspring age | -0.02 | -0.04 | 0.00 | 1 | 11596 | 8647 |
| $\beta$ Association, no play | -0.15 | -0.19 | -0.11 | 1 | 10987 | 8667 |
| $\beta$ Play day | -0.25 | -0.32 | -0.18 | 1 | 10752 | 8628 |
| $\phi$ | 15.61 | 14.83 | 16.32 | 1 | 10302 | 8146 |
| SD (mother) | 0.24 | 0.16 | 0.33 | 1 | 2501 | 4803 |

Table 9: Posterior summaries for the feeding-proportion model (four-level adjacent-day encoding).

| Parameter | Median | HDI lo | HDI hi | Rhat | ESS bulk | ESS tail |
|---|---|---|---|---|---|---|
| $\beta$ Vegetation greenness (EVI) | -0.03 | -0.05 | 0.00 | 1.000 | 12930 | 9013 |
| $\beta$ Fruit availability | -0.06 | -0.09 | -0.04 | 1.000 | 12749 | 9070 |
| Intercept (count) | 0.12 | -0.03 | 0.25 | 1.001 | 2305 | 3395 |
| $\beta$ Offspring age | 0.02 | -0.01 | 0.05 | 1.000 | 12511 | 9053 |
| $\beta$ Day after play | -0.04 | -0.13 | 0.06 | 1.001 | 13975 | 8357 |
| $\beta$ Day before play | 0.03 | -0.08 | 0.15 | 1.000 | 12776 | 9068 |
| $\beta$ Day of play | -0.23 | -0.30 | -0.16 | 1.000 | 14061 | 8373 |
| $\phi$ | 16.06 | 15.01 | 17.15 | 1.000 | 15003 | 8910 |
| SD (mother) | 0.31 | 0.20 | 0.43 | 1.000 | 2537 | 5335 |

Table 10: Posterior summaries for the Approach model. $\beta$ Day before play and Within-day before play are the categorical phase contrasts vs. baseline.

| Parameter | Median | HDI lo | HDI hi | Rhat | ESS bulk | ESS tail |
|---|---|---|---|---|---|---|
| $\beta$ Daily path length | 0.00 | -0.01 | 0.01 | 1 | 16410 | 9092 |
| $\beta$ Vegetation greenness (EVI) | 0.00 | -0.01 | 0.01 | 1 | 19454 | 8408 |
| $\beta$ Fruit availability | 0.00 | -0.02 | 0.01 | 1 | 15434 | 9462 |
| Intercept (count) | 0.00 | -0.02 | 0.01 | 1 | 8088 | 7821 |
| $\beta$ Mode separation | 0.00 | -0.01 | 0.01 | 1 | 12720 | 9436 |
| $\beta$ Offspring age difference | 0.00 | -0.01 | 0.01 | 1 | 17059 | 9173 |
| $\beta$ Offspring age (sum) | 0.01 | 0.00 | 0.02 | 1 | 17107 | 9523 |
| $\beta$ Day before play | 0.09 | 0.03 | 0.16 | 1 | 16122 | 8416 |
| $\beta$ Within-day before play | -0.01 | -0.11 | 0.09 | 1 | 8823 | 8506 |
| $\beta$ Relatedness | 0.00 | -0.01 | 0.01 | 1 | 11281 | 9563 |
| $\beta$ Window duration | -0.02 | -0.04 | 0.00 | 1 | 9156 | 8890 |
| SD (dyad) | 0.01 | 0.00 | 0.02 | 1 | 4475 | 5051 |
| SD (mother) | 0.01 | 0.00 | 0.03 | 1 | 4556 | 5692 |
| $\sigma$ | 0.28 | 0.27 | 0.29 | 1 | 17693 | 8376 |

Table 11: Posterior summaries for the Return model. $\beta$ Within-day after play and Day after play are the categorical phase contrasts vs. baseline.

| Parameter | Median | HDI lo | HDI hi | Rhat | ESS bulk | ESS tail |
|---|---|---|---|---|---|---|
| $\beta$ Daily path length | -0.01 | -0.02 | 0.00 | 1 | 17671 | 8818 |
| $\beta$ Vegetation greenness (EVI) | 0.00 | -0.01 | 0.01 | 1 | 19822 | 9040 |
| $\beta$ Fruit availability | 0.00 | -0.01 | 0.01 | 1 | 17209 | 9494 |
| Intercept (count) | -0.02 | -0.04 | 0.00 | 1 | 5736 | 6609 |
| $\beta$ Mode separation | 0.00 | -0.01 | 0.01 | 1 | 15708 | 9395 |
| $\beta$ Offspring age difference | 0.00 | -0.01 | 0.01 | 1 | 19869 | 9135 |
| $\beta$ Offspring age (sum) | 0.00 | -0.01 | 0.01 | 1 | 18756 | 9985 |
| $\beta$ Day after play | -0.01 | -0.05 | 0.04 | 1 | 16940 | 9008 |
| $\beta$ Within-day after play | 0.07 | 0.03 | 0.12 | 1 | 12165 | 9511 |
| $\beta$ Relatedness | 0.00 | -0.01 | 0.01 | 1 | 11530 | 8957 |
| $\beta$ Window duration | 0.01 | 0.00 | 0.02 | 1 | 13369 | 9792 |
| SD (dyad) | 0.01 | 0.00 | 0.02 | 1 | 6311 | 6145 |
| SD (mother) | 0.02 | 0.01 | 0.05 | 1 | 4533 | 4792 |
| $\sigma$ | 0.25 | 0.24 | 0.25 | 1 | 18022 | 8518 |
